# PRMT5 inhibition induces a histone methylation switch that drives chromatin plasticity, drug resistance, and metastatic progression

**DOI:** 10.64898/2026.01.30.702866

**Authors:** Colin E. Fowler, Natalie A. O’Hearn, Nicole Henning, Anthony Griffen, Ziyang Ye, Nicolas Mathey-Andrews, Griffin J. Salus, Renin Hazan, Tyler Jacks, Aurora A. Burds, Helen S. Mueller, Yadira Soto-Feliciano, Lindsay M. LaFave, Jacqueline A. Lees

## Abstract

PRMT5 is the major type II protein arginine methyltransferase, regulating fundamental processes including transcription, splicing, and translation. Its expression is restricted to stem and progenitor cells but reactivated in many tumors. Numerous PRMT5 inhibitors (PRMT5i) have been developed, and several are currently in clinical trials. Their efficacy is largely attributed to disruption of PRMT5’s role in splicing. Here we show that PRMT5i produce dichotomous effects in lung adenocarcinoma (LUAD): they induce lethal splicing defects but also trigger epigenetic changes that actively enable acquisition of drug resistance. Of concern, the resulting stably resistant populations have undergone accelerated dedifferentiation, gaining molecular and phenotypic hallmarks of metastatic disease. Moreover, even brief PRMT5i treatment of drug-naïve tumors is sufficient to drive disease progression *in vivo*. This epigenetic rewiring reflects a previously unappreciated dominance of PRMT5 over PRMT1, arising from differential subcellular localization. Our data indicate that cytoplasmic PRMT5 likely dimethylates histones symmetrically before nuclear import. PRMT5i block this, allowing unmodified histones to reach nuclear PRMT1 and undergo asymmetric dimethylation instead. This symmetric-to-asymmetric switch flips histones from repressive to activating states, yielding rapid chromatin opening and derepression of lineage programs, a subset of which are selected for and retained in resistant cells. Accordingly, inhibition of asymmetric dimethylation suppresses PRMT5i-induced chromatin derepression and synergizes strongly with PRMT5 inhibition. Finally, endogenous PRMT5 suppression in *MTAP*-deficient human cells partially engages this switch, and *CDKN2A/MTAP*-deficient tumors are correspondingly more dedifferentiated. Thus, PRMT5i create a splicing vulnerability while simultaneously licensing epigenetic escape toward a more dedifferentiated and aggressive cell state.

**Significance Statement:** PRMT5i can drive lung cancer progression by inducing a previously unrecognized histone methylation switch that promotes drug resistance and dedifferentiation, raising concern about clinical deployment of these inhibitors.

## Introduction

Arginine methylation is a common post-translational modification, catalyzed by members of the protein arginine methyltransferase (PRMT) family, that enables protein-protein interactions^1^. Type I PRMTs catalyze asymmetric dimethylarginine (ADMA), while type II PRMTs catalyze symmetric dimethylarginine (SDMA). PRMT5 is the major type II PRMT and many of its substrates regulate fundamental molecular processes, including transcription, splicing, and translation^2^. Early studies established PRMT5 as a critical regulator of development and cell identity^3,4^. PRMT5 is highly expressed in embryonic stem (ES) cells and is downregulated as cells differentiate^5^. In ES cells, PRMT5 maintains lineage fidelity through repression of developmental gene expression programs, in part through symmetric dimethylation of histone H2A arginine 3 (H2AR3me2*s*)^3^. Loss of PRMT5 leads to derepression of developmental programs, profound defects in cell fate specification, and embryonic lethality^3^, underscoring its essential role in maintaining stable epigenetic states.

PRMT5 also plays a central role in RNA splicing, functioning in the methylosome complex to catalyze SDMA modifications of spliceosomal Sm proteins^6–11^. Disruption of PRMT5 activity causes widespread splicing defects, most prominently the accumulation of detained introns (DIs)^5,12,13^ – intronic sequences retained in otherwise fully processed transcripts, which remain trapped in the nucleus^14^. This nuclear sequestration reduces levels of cytoplasmic transcripts available for translation^5,12,13^. Interestingly, PRMT5-regulated DI-containing genes are strongly enriched in regulators of cell cycle progression and DNA damage response^5,12,13^. Accordingly, disruption of PRMT5 activity induces cell cycle defects, DNA damage signatures, and consequent cell death^5,12,13^.

PRMT5 is lowly expressed in terminally differentiated tissues, but shows near-universal re-expression in cancer^2,15,16^. Importantly, elevated expression is associated with poor clinical outcomes^15–17^, and many cancer cells require PRMT5 for viability. Accordingly, numerous PRMT5 inhibitors (PRMT5i) have been developed, several of which are advancing into early-stage clinical trials^18,19^. PRMT5’s role in splicing, and particularly in suppression of DI formation, has been established as the primary driver of therapeutic sensitivity^5,12,13,20–23^. Thus, loss of Sm protein methylation and the appearance of splicing defects are key pharmacodynamic biomarkers of PRMT5 inhibition^20,23–27^.

Even though multiple PRMT5i have entered clinical trials, little is known about the durability of response and the potential for resistance. We have previously shown that stable PRMT5i resistance arises rapidly in LUAD cell lines derived from the autochthonous *Kras*^G12D/+^;*Trp53*^fl/fl^ (KP-mutant) model and reflects drug-induced changes rather than selection of a pre-existing state^28^. KP-mutant LUAD tumors progress along a well-defined dedifferentiation continuum, losing lung identity (including decreased NKX2-1 expression) as they move through a gastric-marked transition (HNF4A) toward an increasingly embryonic-like character (HMGA2)^29–31^. These transitions are driven by coordinated transcriptional and epigenomic remodeling^31–33^, and dedifferentiation is similarly associated with disease progression and worse outcomes in human LUAD^34–36^.

Here, we investigate the mechanisms and consequences of acquired resistance to PRMT5 inhibition in KP-mutant LUAD. We find that resistant cells do not restore splicing fidelity, as they display the same splicing defects as parental controls in response to PRMT5i treatment. Instead, resistant cells shift stably to a late-stage phenotype along the normal KP-mutant LUAD dedifferentiation trajectory. Accordingly, the resistant cells display increased metastatic behavior *in vivo*. Our data further argue that short-term PRMT5 inhibition actively drives this state shift, effectively enabling its own resistance. Specifically, PRMT5i induces widespread chromatin opening and derepression of lineage programs, generating epigenetic plasticity from which resistant states emerge and are subsequently selected. Mechanistically, our data reveal a previously unappreciated interplay between PRMT5 and PRMT1 that enhances the switch from transcriptional repression to activation upon PRMT5 inhibition. This is driven by the differential subcellular localization of these two enzymes and their competing modification of H2AR3. In the basal context, PRMT5 is dominant over PRMT1 because its cytoplasmic localization allows it to catalyze the repressive H2AR3me2*s* mark. When PRMT5 is inhibited, H2A enters the nucleus in an unmethylated form, where it encounters the nuclear-localized PRMT1 and undergoes asymmetric dimethylation to yield a large pool of the active asymmetric dimethylation of H2AR3 (H2AR3me2*a*). Finally, we show that this drug-induced state shift is not confined to our mouse model: human *CDKN2A*/*MTAP*-deficient LUAD and melanoma tumors, which harbor endogenous suppression of PRMT5 activity, display a similarly elevated degree of dedifferentiation. Collectively, our data show that PRMT5 inhibition has dichotomous effects on cancer cells: it creates a vulnerability through splicing defects while simultaneously licensing drug escape by unlocking chromatin plasticity and enabling tumor evolution.

## Results

### PRMT5i-induced splicing defects are not rescued in PRMT5i-resistant lines

We previously generated numerous variants of KP-mutant LUAD cell lines that acquired stable resistance to PRMT5i (**Fig. 1A** and **Fig. S1A**)^28^. To investigate the molecular events that underlie resistance, we compared KP^Early^- and KP^Late^-Parental (representing earlier- and later-stage points along the KP-mutant LUAD dedifferentiation continuum, respectively) with representative PRMT5i-resistant counterparts, hereafter KP^Early^-Resistant and KP^Late^-Resistant (line R1 of three independently derived resistant lines per parental line). Although these lines were selected in EPZ015666, they were also cross-resistant to the mechanistically distinct, S-adenosylmethionine (SAM)-competitive inhibitor JNJ-64619178 (**Fig. 1B** and **Fig. S1B**)^28^. Because PRMT5i lethality is driven by the induction of DIs^12^, we reasoned that resistant cells would have blunted the splicing response to PRMT5i. To test this, we treated our paired cell lines with two mechanistically distinct PRMT5i, the SAM-competitor JNJ-64619178 (for KP^Early^) and substrate-competitor EPZ015666 (for KP^Late^). We used drug doses that maximized the differential viability of parental versus resistant lines and limited treatment to 3 days, as we have previously shown that this yields splicing defects in the parental lines but precedes onset of phenotypic defects^12^. Poly(A)+ mRNA was isolated and deep sequenced (>40 million reads per sample), followed by analysis and quantification of alternative splicing (AS) events. Consistent with our prior studies^5,12^, PRMT5i induced numerous AS events in the parental lines, the vast majority of which involved increased DI formation (**Fig. 1C** and **Fig. S1C**), and gene ontology (GO)^37,38^ analyses confirmed that genes harboring PRMT5i-induced DIs were significantly enriched for proliferation-associated processes (**Fig. S1D**). Notably, contrary to our hypothesis, splicing alterations were remarkably similar between parental and resistant cells despite their differential sensitivity to PRMT5i. First, the identities of DIs significantly upregulated by PRMT5i treatment were highly concordant between parental and resistant lines (p<10^−213^ for KP^Early^ and p<10^−221^ for KP^Late^; **Fig. 1D** and **Fig. S1E–F**), as were the much rarer exon-based AS events (p<10^−127^ for KP^Early^ and p<10^−153^ for KP^Late^; **Fig. S1F**). Second, the magnitude of DI induction (as well as the exon-based splicing changes) was strongly correlated between parental and resistant cells, with Deming regression indicating nearly identical responses (**Fig. 1E** and **Fig. S1G**). Quantitative real-time reverse transcription PCR (qRT-PCR) analysis of representative PRMT5i-responsive DIs (in *Sf3b2* and *Sos2*) confirmed induction upon treatment (p<0.001), with comparable magnitudes in parental and resistant cells (**Fig. S1H**).

**Figure 1.**
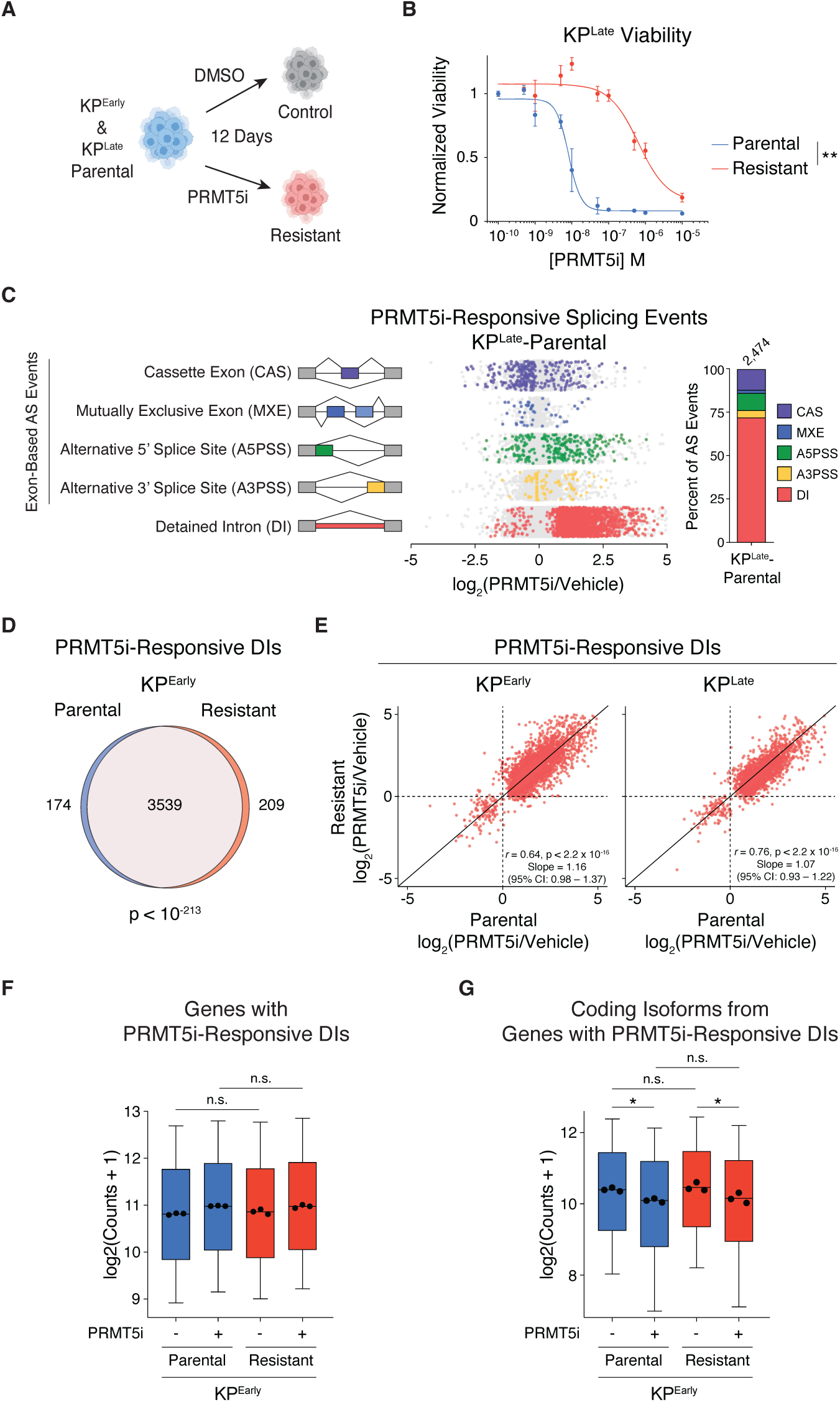
PRMT5i induces equivalent DI formation in sensitive and resistant cells. **A.** Schematic of method used to generate control (DMSO) and resistant (10 μM EPZ015666) variants of KP^Early^ and KP^Late^ LUAD cell lines^28^. **B.** Relative viability of KP^Late^-Parental (blue) versus KP^Late^-Resistant (red), after a 5-day treatment with JNJ-64619178 (PRMT5i). Data are mean ± SD of 3 technical replicates per line. Significance compares area under the curve (AUC) by Welch’s t-test. **C.** KP^Late^-Parental cells were treated with vehicle or 10 μM EPZ015666 (PRMT5i) for 3 days and poly(A)+ mRNA deep sequenced to identify and quantify DIs, as well as exon-based alternative splicing (AS) events: cassette exons (CAS), mutually exclusive exons (MXE), and alternative 5′ or 3′ splice sites (A5PSS, A3PSS). (*Left*) Dots represent individual splicing events and denote those significantly (p_adj_<0.05, color) or not significantly (gray) altered by PRMT5i. (*Right*) Graph shows the proportion of each splicing type that is significantly (p_adj_<0.05) altered by PRMT5i, with the total number of significant events indicated above. **D.** Overlap of DIs significantly altered (PRMT5i-responsive) after 3-day treatment of KP^Early^-Parental versus KP^Early^-Resistant with 100 nM JNJ-64619178. Significance determined by hypergeometric test. **E–G.** Parental and resistant lines were treated for 3 days with vehicle or PRMT5i (10 μM EPZ015666 for KP^Late^-Parental and KP^Late^-Resistant or 100 nM JNJ-64619178 for KP^Early^-Parental and KP^Early^-Resistant). **E.** Graphs compare the induction levels of PRMT5i-responsive DIs (p_adj_<0.05) in parental versus resistant lines for KP^Early^ (*Left*) and KP^Late^ (*Right*). Dots represent individual DIs and the solid line indicates a line of identity. Pearson correlation and Deming regression values are displayed. **F–G.** Graphs compare the normalized expression counts of: (**F**) genes with PRMT5i-responsive DIs or (**G**) their fully processed (i.e. non-DI containing) coding isoforms, for KP^Early^-Parental (blue) and KP^Early^-Resistant (red) with vehicle (-) or PRMT5i (+) treatment. Box plots were generated from average log_2_(Counts+1) for each gene (n=2,257), with individual points overlaid representing median counts for each replicate (n=3). Two-way ANOVA, with treatment and resistance status as main effects, was performed on the median counts for each replicate. For all panels, * indicates p<0.05, ** indicates p<0.01, and n.s. indicates not significant.

DIs reduce gene expression by retaining transcripts in the nucleus, thereby decreasing the levels of fully processed cytoplasmic mRNAs available for translation^5,12,14^. Given the persistence of splicing defects in resistant cells, we wondered whether these cells display increased expression of genes that harbor PRMT5i-induced DIs, either before or in response to PRMT5i treatment. To address this, we considered genes in which PRMT5i significantly alters DI formation and quantified the levels of total transcripts, as well as the levels of DI-containing isoforms versus fully spliced, coding isoforms using RSEM^39^ (**Fig. S2A**). For both KP^Early^ and KP^Late^, we saw no significant differences in the total levels of DI-containing genes between resistant and parental lines at baseline and no consistent differences following PRMT5 inhibition (**Fig. 1F** and **Fig. S2B**). Moreover, in resistant versus parental lines, PRMT5i yielded comparable increases in the levels of DI-containing isoforms (**Fig. S2C**) and comparable, significant (p<0.05) reductions in the levels of the properly spliced coding isoforms (**Fig. 1G** and **Fig. S2D**). Collectively, these findings establish that our PRMT5i-resistant cells neither restore splicing fidelity nor compensate by increasing abundance of affected genes. Given this outcome, we hypothesized that resistance results from adoption of a distinct cellular state, which enables tolerance of the induced splicing defects.

### PRMT5i resistance drives a stable shift toward advanced, dedifferentiated KP-mutant LUAD states

Cellular plasticity frequently accompanies both tumor progression and the acquisition of drug resistance. Progression of KP-mutant tumors follows a well-defined dedifferentiation continuum that is marked by sequential changes in lineage-defining programs, essentially reversing normal lung development (**Fig. 2A**). Our parental lines were chosen for this study because they reside at different points on the KP-mutant LUAD dedifferentiation continuum (as indicated by their names, KP^Early^ and KP^Late^). We previously showed that PRMT5i-resistant cells possess distinct transcriptional signatures from their parental controls^28^. Given this, we wondered whether they remain on the dedifferentiation continuum or have instead entered an alternate lineage state. To address this, we performed differential gene expression analysis on RNA-Seq data from KP^Early^- and KP^Late^-Parental cells versus biological triplicate samples for three independently derived resistant populations for each cell line (KP^Early^-Resistant 1, 2, and 3 and KP^Late^-Resistant 1, 2, and 3; **Fig. S1A**). Principal component analysis (PCA) revealed that the resistant populations all segregate from their parental counterparts, confirming that resistance is associated with transcriptional state transitions (**Fig. 2B** and **Fig. S3A**). Gene set enrichment analysis (GSEA)^40^ established that the resistant lines showed coordinated activation of transcriptional programs associated with advanced KP-mutant LUAD progression, including epithelial-to-mesenchymal transition (EMT), gastric lineage programs, and metastasis (**Fig. 2C** and **Fig. S3B**). Matching these pathway-level changes, we observed major changes in RNA and protein levels for canonical KP-mutant LUAD differentiation markers (**Fig. 2D–E**). For KP^Early^, the parental cells displayed a lung lineage-like profile, characterized by high expression of the lung marker *Nkx2-1*. The three KP^Early^-Resistant derivatives all lost *Nkx2-1* expression, downregulated *Hnf4a* at the RNA level, and upregulated *Hmga2* at both the RNA and protein levels, indicating progression along the dedifferentiation continuum past the gastric-associated transition and toward an increasingly embryonic-like character. KP^Late^ cells, which originate from more advanced disease (indicated by the absence of *Nkx2-1* expression in parental cells), also gained notable dedifferentiation in the resistant variants, marked by profound downregulation of *Hnf4a* and induction of *Hmga2* at both the RNA and protein levels, consistent with progression further along that same continuum. Although KP-mutant LUAD can transdifferentiate into squamous or neuroendocrine states under therapeutic pressure^41–46^, markers of these alternative lineages were not induced in our resistant cells (**Fig. S3C**). Together, these data indicate that PRMT5i resistance reflects a shift along the dedifferentiation continuum toward advanced disease, regardless of the starting differentiation state.

**Figure 2.**
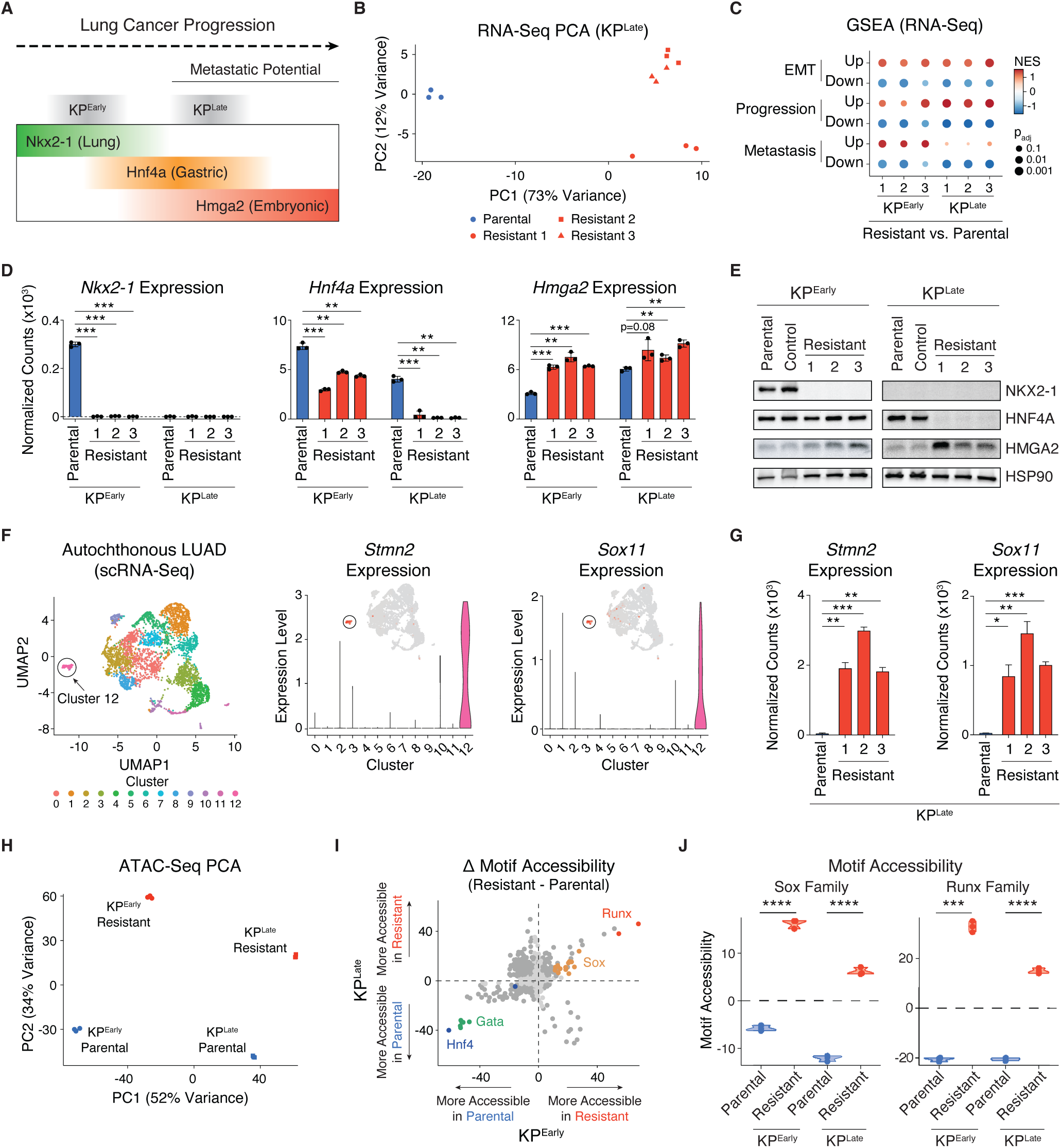
PRMT5i-resistant KP-mutant LUAD cells exhibit transcriptional and chromatin accessibility signatures indicative of more progressed disease. **A.** Schematic of KP-mutant LUAD dedifferentiation spectrum, including the relative states of KP^Early^- and KP^Late^-Parental lines. Each bar represents the general expression pattern of the given dedifferentiation state marker. **B.** PCA of the 500 most variably expressed genes by RNA-Seq in KP^Late^-Parental (blue) versus three independently derived KP^Late^-Resistant populations (red) with triplicate samples for each. **C.** GSEA results for gene sets listed in **Fig. S3B**, drawn from published signatures representing genes increased or decreased during each indicated process, comparing KP^Early^- and KP^Late^-Parental to Resistant lines. Color represents normalized enrichment score and point size is scaled inversely to FDR, such that larger points indicate greater significance. **D.** Relative expression of dedifferentiation markers (*Nkx2-1*, *Hnf4a*, *Hmga2*) in KP^Early^- and KP^Late^-Parental (blue) and Resistant (red) lines. Data are mean ± SD of 3 biological replicates per line. **E.** Western blot of core KP-mutant LUAD dedifferentiation markers (NKX2-1, HNF4A, HMGA2) and HSP90 loading control in KP^Early^- and KP^Late^-Parental, Control (DMSO selection), and Resistant lines. **F.** UMAP of scRNA-Seq dataset from an autochthonous model of KP-mutant LUAD with (*left*) all clusters marked, (*center*) *Stmn2* expression, or (*right*) *Sox11* expression within individual clusters. Cluster 12 (circled) was previously described to contain late-stage, pro-metastatic cells^32^. **G.** Normalized RNA expression counts for *Stmn2* and *Sox11* in KP^Late^-Parental (blue) and Resistant (red) lines. Data are mean ± SD of 3 biological replicates per line. **H.** PCA of the 5,000 most variable ATAC-Seq peaks across KP^Early^- and KP^Late^-Parental and Resistant lines, with three biological replicates per line. KP^Early^ samples are shown as circles and KP^Late^ samples as squares; parental lines are shown in blue and resistant lines in red. **I.** ΔchromVAR deviation scores (Resistant–Parental) for KP^Early^ and KP^Late^, reflecting relative changes in TF motif accessibility between resistant and parental lines. Dark gray or colored points denote motifs that show significant differential accessibility in both comparisons (p_adj_<0.01), while light gray points are not significant. Colored points indicate motifs from the same TF family. **J.** chromVAR deviation scores for combined (*left*) Sox and (*right*) Runx family motifs in KP^Early^- and KP^Late^-Parental (blue) and Resistant (red) lines. Each point represents a biological replicate (n=3). For corresponding panels, Welch’s t-test was used. * indicates p<0.05, ** indicates p<0.01, *** indicates p<0.001, and **** indicates p<0.0001.

To determine the extent of progression, we took advantage of a single-cell RNA-Seq dataset that defined a continuum of tumor states from autochthonous KP-mutant LUAD^32^. This culminates in a small population of cells (Cluster 12) that are highly dedifferentiated (**Fig. 2F**). In addition to high *Hmga2* expression (**Fig. S3D**), Cluster 12 is distinguished from all other cell populations by high expression of *Sox11* and *Stmn2* (**Fig. 2F**). Strikingly, both of these markers are implicated in cancer^47–53^ and strongly induced in KP^Late^-Resistant cells (**Fig. 2G**). Moreover, we previously found that *Stmn2* is functionally required for KP^Late^ cells to acquire PRMT5i-resistance^28^. Thus, PRMT5i-resistant cells display transcriptional features associated with the most advanced stages of KP-mutant LUAD.

To determine whether the resistant transcriptional state is underpinned by stable epigenetic remodeling, we performed the Assay for Transposase-Accessible Chromatin with sequencing (ATAC-Seq) on KP^Early^- and KP^Late^-Parental and Resistant lines. The identified chromatin accessibility changes closely mirrored transcriptional alterations at key marker loci, indicating coordinated regulatory reprogramming. For example, accessibility was markedly reduced at the *Nkx2-1* locus in KP^Early^-Resistant cells and at *Hnf4a* in both KP^Early^- and KP^Late^-Resistant cells (**Fig. S3E**), mirroring their progression along the dedifferentiation continuum. Accordingly, genome-wide analyses revealed that chromatin accessibility profiles robustly distinguished resistant from parental cells. PCA separated samples by both disease stage (PC1) and resistance status (PC2), indicating that resistance is associated with a conserved accessibility shift across independent models (**Fig. 2H**). Differential accessibility analysis identified thousands of regions that were gained or lost in resistant cells relative to their parental counterparts (**Fig. S3F**), which we define as resistant state loci. Importantly, many of these accessibility changes were shared between the resistant populations of both KP^Early^ and KP^Late^ (**Fig. S3G**), supporting the existence of a common resistance-associated chromatin state.

To define the regulatory features of this state, we analyzed transcription factor (TF) motif accessibility using chromVAR^54^. Resistant cells exhibited reduced accessibility at motifs associated with lineage-specifying factors, including Nkx and Gata family motifs, which include established regulators of lung identity (e.g., NKX2-1, GATA6), and Hnf4 family motifs, including HNF4A, a marker of the gastric-associated transition along the KP-mutant LUAD dedifferentiation continuum, alongside increased accessibility at motifs linked to lineage plasticity, metastasis, and advanced disease states, particularly Sox and Runx family motifs (**Fig. 2I–J** and **Fig. S3H**), which are known regulators of metastasis^31,48,55–58^. Notably, these motif accessibility changes were highly concordant between KP^Early^ and KP^Late^ lines (**Fig. 2I**), consistent with the existence of a conserved regulatory state shift accompanying resistance acquisition. Collectively, these data demonstrate that PRMT5i resistance is associated with stable epigenetic reprogramming characterized by loss of lineage-maintenance networks and activation of regulatory programs linked to lineage plasticity and advanced disease.

### PRMT5i-resistant tumors are hyper-metastatic, and PRMT5 inhibition itself can drive progression

Having established that PRMT5i-resistant cells possess transcriptional and chromatin signatures characteristic of advanced KP-mutant LUAD, we directly tested their metastatic potential using multiple *in vivo* models, as compared to parental controls (**Fig. S4A–B**). We first evaluated the metastatic potential of KP^Late^-Parental versus Resistant cells (1×10^5^ cells, 5–7 recipients per line) by tail vein injection. While yielding comparable lung tumor burden (**Fig. 3A**), the KP^Late^-Resistant tumors were significantly more invasive, generating more pleural invasions (p<0.05) as well as liver metastases that were both more frequent (p<0.05) and substantially larger (p<0.01; **Fig. 3A–B**). Since tail vein injections bias toward pulmonary colonization, we next used ultrasound-guided injection (USGI) into the left ventricle^59^, which enables systemic distribution of cells to a broad spectrum of peripheral organs (**Fig. S4A**). Following intracardiac injection (5×10^3^ cells, 7–8 recipients per line), KP^Late^-Parental and Resistant cells again yielded lung tumors of comparable burden, while the KP^Late^-Resistant cells again produced significantly more pleural invasions (p<0.01) and formed larger, more aggressive liver metastases (p<0.05; **Fig. S4C–D**). We also tested KP^Early^ cells in this assay and found that the resistant variant yielded increased lung tumor burden, as well as a markedly higher frequency and size of liver metastases, compared to parental controls (**Fig. S4E–G**).

**Figure 3.**
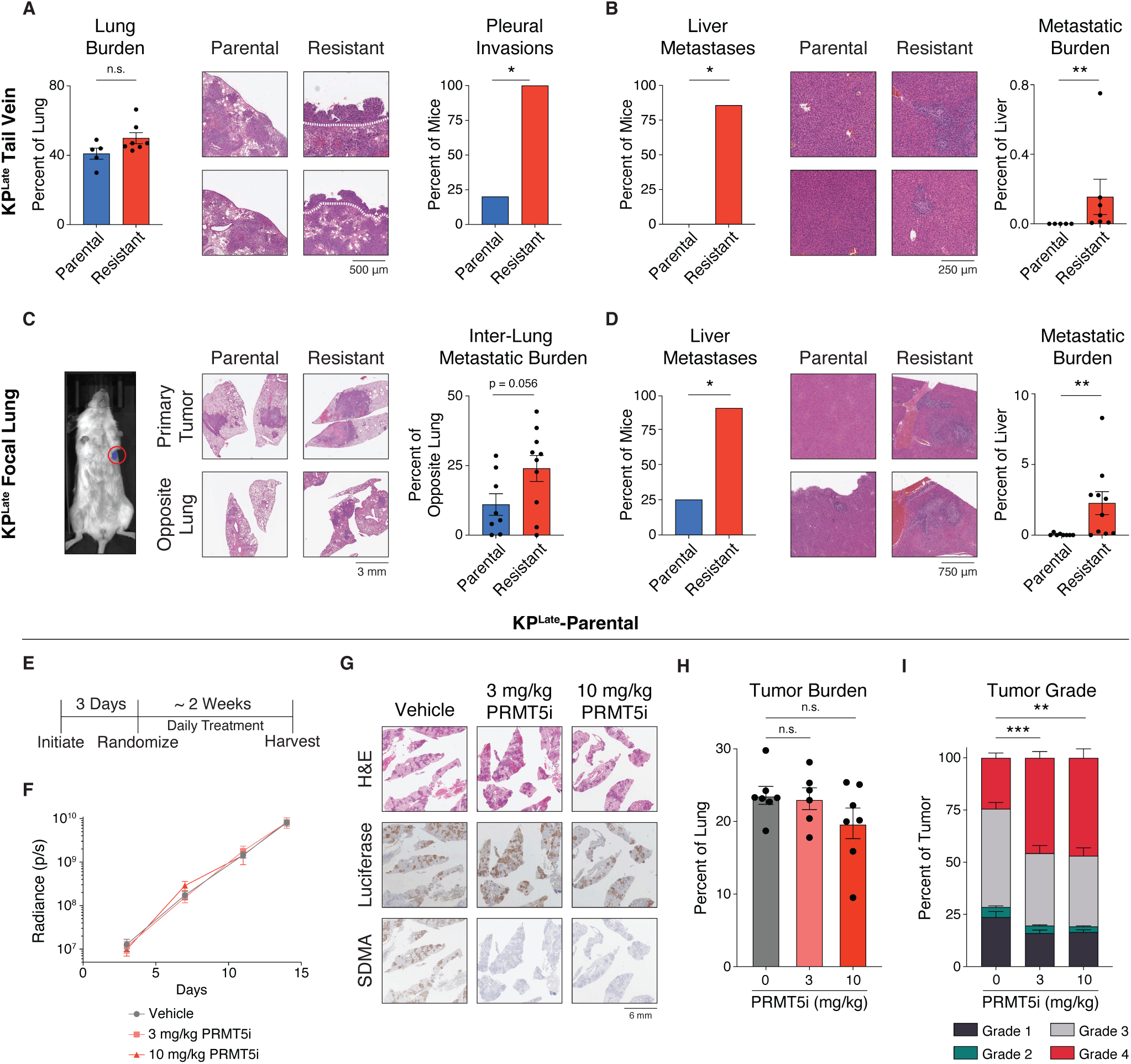
PRMT5i-resistant cells have increased metastatic potential *in vivo* compared to their parental lines. **A–B.** Tail vein injection was used to introduce 1×10^5^ KP^Late^-Parental or Resistant cells into 5–7 recipients per line. These were analyzed for: **(A)** lung tumors, with tumor burden (p=0.11), representative H&E images, and pleural invasion quantification (p=0.010) shown; and **(B)** liver metastases, with metastatic frequency (p=0.015), representative H&E images, and burden (p=0.0025) shown. **C–D.** Focal lung ultrasound-guided injection (USGI) was used to introduce 5×10^3^ luciferase-expressing KP^Late^-Parental or Resistant cells into 8–10 recipients per line. These were analyzed for: **(C)** lung tumors, with representative IVIS and H&E images, and inter-lung metastatic burden (p=0.056) shown; and **(D)** liver metastases, with metastatic frequency (p=0.013), representative H&E images, and burden (p=0.0017) shown. **E–I.** As schematized in (**E**), tail vein injection was used to introduce 5×10^3^ luciferase-expressing KP^Late^-Parental cells into 6–7 recipients per condition. Mice were randomized into cohorts receiving 0 (vehicle), 3, or 10 mg/kg PRMT5i daily and analyzed for: (**F**) relative luminescence over time; (**G**) representative lung histology sections including H&E, luciferase IHC, and SDMA IHC; (**H**) lung tumor burden (p=0.82 for 3 mg/kg and p=0.15 for 10 mg/kg compared to vehicle); and (**I**) lung tumor grade, where for Grade 4 tumor percentage p=0.0003 for 3 mg/kg and p=0.0014 for 10 mg/kg compared to vehicle. For all panels, data are mean ± SEM where applicable. Fisher’s exact test was used for frequency of tumors, metastases, and pleural invasions, Mann-Whitney U-test was used for burden, and Welch’s t-test was used for all *in vivo* treatment burden and grade. * indicates p<0.05, ** indicates p<0.01, *** indicates p<0.001, and n.s. indicates not significant.

Metastasis is a multistep process involving escape from the primary tumor, intravasation, circulation, extravasation, and colonization; however, tail vein and intracardiac models bypass the first two steps of this cascade. To assess the complete metastatic cascade, we performed focal left lung USGI (**Fig. S4A**) with KP^Late^-Parental and Resistant cells (5×10^3^ cells, 8–10 recipients per line). Intravital bioluminescence (IVIS) imaging and histological analysis confirmed formation of single primary lung tumors in all recipients (**Fig. 3C**). We then assessed metastasis to the contralateral lung and the liver. KP^Late^-Resistant tumors exhibited increased inter-lung metastatic burden (p=0.056) and significantly more frequent and larger liver metastases (p<0.05 and p<0.01, respectively; **Fig. 3C–D**). Metastases arising from resistant cells were highly invasive, infiltrating surrounding tissue and establishing tumors within liver vasculature. Thus, PRMT5i-resistant cells efficiently execute the full metastatic cascade and display heightened aggressiveness *in vivo*.

Given these striking phenotypes, we next asked whether PRMT5i treatment is able to drive disease progression entirely in the *in vivo* setting. For this, we seeded lung tumors via tail vein injection of luciferase-expressing KP^Late^-Parental cells (5×10^3^ cells), confirmed the presence of tumor burden by IVIS imaging, and then randomized the recipients into treatment cohorts receiving 0 (vehicle), 3, or 10 mg/kg PRMT5 inhibitor daily (n=6–7 per cohort; **Fig. 3E**). We quantified lung tumor growth at periodic timepoints by IVIS imaging, as well as terminal tumor burden from histological sections. We found that PRMT5i treatment yielded no significant reductions in either the rate of tumor growth or terminal burden (**Fig. 3F–H**), even though both drug doses successfully inhibited PRMT5 activity, as indicated by complete loss of immunohistochemical staining for SDMA (**Fig. 3G**). Thus, effective PRMT5 inhibition failed to reduce the burden of established lung tumors. We then assessed disease progression by tumor grade analysis and found that both PRMT5i treatment doses induced a pronounced shift toward higher-grade lesions, with grade 4 tumors increasing nearly two-fold and corresponding decreases in lower-grade tumor frequencies (**Fig. 3I** and **Fig. S4H**). Together, these data demonstrate that the molecular rewiring associated with PRMT5i resistance – including dedifferentiation, transcriptional remodeling, and chromatin accessibility changes – not only confers survival advantage in the presence of PRMT5i but also drives aggressive metastatic behavior. Moreover, pharmacological inhibition of PRMT5 accelerates disease progression *in vivo* without any accompanying reduction in lung tumor burden.

### PRMT5i treatment derepresses chromatin to enable cellular plasticity and resistance

Our transcriptional, chromatin accessibility, and metastatic phenotype data support the notion that PRMT5i resistance involves a shift to a distinct epigenetic state. We and others have established that PRMT5’s splicing function underlies the vulnerability of cancer cells to PRMT5 inhibition^5,60–63^. However, PRMT5 has another key function in normal development in which it catalyzes SDMA modifications of histones that promote chromatin compaction and repress differentiation-associated loci^3,4^. The existence of this mechanism, together with our prior discovery that PRMT5i resistance is drug-induced, not pre-existing^28^, suggested that PRMT5i treatment might itself drive chromatin opening to yield widespread derepression of lineage-specification programs. This could drive cellular heterogeneity, enabling selection of the epigenetic state(s) that provide resistance.

To test the first part of our hypothesis, we asked whether short-term exposure to PRMT5i impacts global chromatin accessibility. For this, we used ATAC-See^64^, in which Tn5 transposase bearing fluorescently labeled adapters inserts preferentially into regions of open chromatin, allowing for single-cell quantification of chromatin accessibility by flow cytometry (**Fig. 4A**). We applied this approach to KP^Early^- and KP^Late^-Parental cells treated with vehicle or PRMT5i for 5 days, a timepoint at which both lines exhibit profound loss of SDMA (**Fig. S5A**). Since PRMT5i treatment alters cell cycle phasing (**Fig. S5B**), we examined the ATAC-See signals of G1 and G2 populations separately (**Fig. 4B** and **Fig. S5C**). In both phases, the PRMT5i-treated samples had significantly higher signals for KP^Early^- and KP^Late^-Parental (p<0.01; **Fig. 4B**), indicating broad chromatin opening. To identify the affected loci, we performed bulk ATAC-Seq at the same 5-day timepoint. Differential chromatin accessibility analysis between vehicle versus drug treated samples (p_adj_<0.01, |log_2_FC|>1; **Fig. 4C** and **Fig. S5D**) showed that short-term PRMT5i treatment opens substantially more loci than it closes (>2.2-fold), consistent with PRMT5’s role as a chromatin repressor^3,65,66^. If PRMT5i treatment enables the creation and sampling of a heterogeneous mix of cells with alternative chromatin states, we reasoned that loci gaining accessibility specifically under PRMT5i should show greater replicate-to-replicate variability than loci gaining accessibility specifically under vehicle. Consistent with this hypothesis, for both KP^Early^- and KP^Late^-Parental lines, PRMT5i-increased loci showed significantly higher coefficient of variation (CV) within PRMT5i replicates than vehicle-increased loci showed within vehicle replicates (p<10^−45^ for KP^Early^-Parental and p<10^−13^ for KP^Late^-Parental; **Fig. 4D** and **Fig. S5E**), supporting the notion that PRMT5i treatment induces chromatin state heterogeneity.

**Figure 4.**
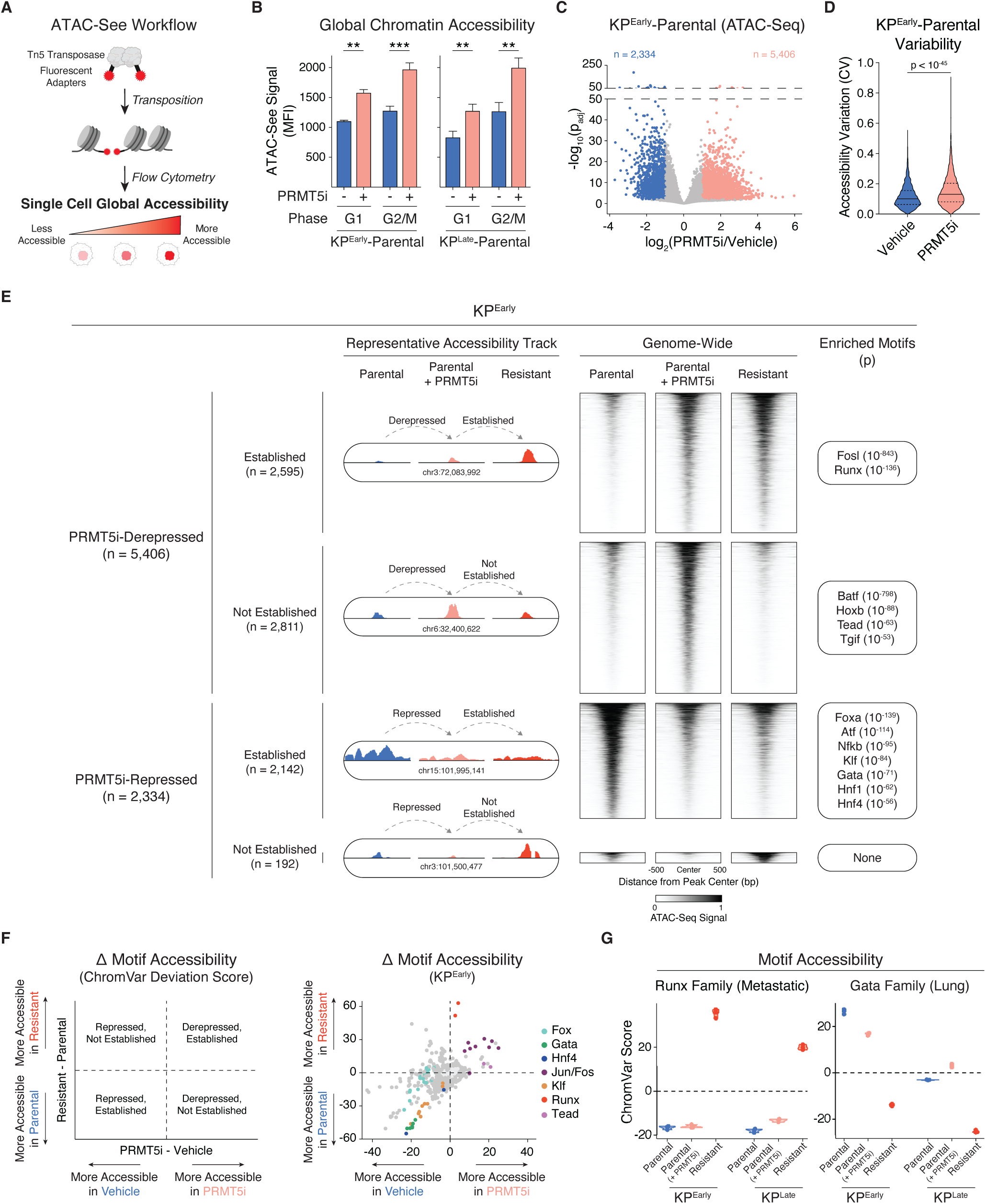
PRMT5 inhibition remodels chromatin accessibility to enable transcriptional plasticity and establishment of resistant state changes. **A.** Schematic of the ATAC-See workflow. **B.** Mean fluorescence intensity (MFI) of ATAC-See signal in cells treated with vehicle or PRMT5i (100 nM JNJ-64619178) for 5 days for both KP^Early^- and KP^Late^-Parental lines. G1 and G2/M populations were considered separately to normalize for DNA content and any effect of drug on cell cycle phasing. Data are mean ± SD of 3 biological replicates per condition. Welch’s t-test. **C.** Differential chromatin accessibility in KP^Early^-Parental cells after 5-day vehicle or PRMT5i treatment. Each point represents an individual genomic locus that is more accessible (pink; p_adj_<0.01, log_2_FC>1), less accessible (blue; p_adj_<0.01, log_2_FC<–1), or not significantly changed (gray). **D.** Coefficient of variation (CV) across replicates for loci with increased accessibility in KP^Early^-Parental cells after 5-day vehicle (blue) or PRMT5i (pink) treatment. Wilcoxon rank-sum test. **E.** Genome-wide analysis of chromatin accessibility changes following 5-day PRMT5 inhibition in KP^Early^-Parental cells and their relationship to accessibility states established in resistant cells. Four categories of loci are shown based on whether PRMT5i induces derepression or repression of accessibility and whether these changes are established or not established in the resistant state: derepressed-and-established, derepressed-and-not established, repressed-and-established, and repressed-and-not established. (*Left*) Representative accessibility tracks. (*Center*) Genome-wide accessibility heatmaps. (*Right*) Enriched *de novo* TF motifs for each category, labeled by best-matching TF (p<10^−50^). **F.** chromVAR was conducted for 5-day vehicle- and PRMT5i-treated KP^Early^-Parental cells (x-axis) and resistant versus parental KP^Early^ cells (y-axis). (*Left*) Schematic illustrating whether motif accessibility changes represent short-term derepression or repression and whether these changes are established or not established in the resistant state. (*Right*) Deviation scores, where individual points represent motifs that are significantly (dark gray or colored, p_adj_<0.01) or non-significantly (light gray) altered. Colored points indicate motifs belonging to the same TF family. **G.** Individual chromVAR deviation scores for selected TF family motifs in KP^Early^ and KP^Late^ showing accessibility in 5-day vehicle (blue) or PRMT5i (pink) treated parental cells and stably resistant cells (red). Each point represents one biological replicate (n=3). For corresponding panels, ** indicates p<0.01 and *** indicates p<0.001.

We then considered the second part of our hypothesis, that the terminal resistant state is selected from the pool of chromatin states induced by short-term PRMT5 inhibition. To address this, we first classified all significant loci changes according to two features: whether they were derepressed or repressed by short-term PRMT5i treatment (“drug-responsive” loci), and whether they were a hallmark of the stably resistant cells (“resistance state” loci; **Fig. 4E–F**). This framework exhaustively partitions drug-responsive loci into four classes that capture both the direction of initial chromatin changes (more accessible/derepressed or less accessible/repressed) and their fates in resistant cells (established or not established). We then used two complementary methods to identify enrichment of TF motifs: Hypergeometric Optimization of Motif EnRichment^67^ (HOMER) to assess patterns within each of the four classes (**Fig. 4E** and **Fig. S6A**) and unbiased chromVAR to analyze global motif accessibility (**Fig. 4F–G** and **Fig. S6B–C**). These two methods yielded concordant results, identifying a relatively small number of significantly altered programs (**Fig. 4E–G** and **Fig. S6B–C**). Importantly, most of these are well-established hallmarks of normal KP-mutant LUAD progression (**Fig. S7A–C**). The Fos/Jun (AP-1) motifs are difficult to interpret, as AP-1 activity is highly context-dependent and can drive divergent transcriptional programs depending on cellular state^68^. AP-1 family members show accessibility within specific segments of KP-mutant LUAD progression (**Fig. S7C**), but also regulate core stress-response genes such that their enrichment could reflect a general stress response to PRMT5 inhibition rather than directed regulatory selection. We therefore focused on the remaining KP-mutant programs. Their behavior was highly consistent with our selection hypothesis for both KP^Early^ and KP^Late^. For example, Runx motifs, which mark progressed metastatic disease^31,57^, were identified as being derepressed-and-established (**Fig. 4E–G** and **Fig. S6A–B**), consistent with selection for late-stage metastatic KP-mutant LUAD programs. At the same time, Tead motifs, which mark the differentiated AT1 cell state^69,70^, were derepressed in response to short-term PRMT5i but were not established in stably resistant cells (**Fig. 4E–G** and **Fig. S6A–C**). This reveals transient accessibility of a more differentiated program that was available but not selected for during resistance acquisition. We also observed key differences in the accessibility dynamics of early lineage programs between KP^Early^ and KP^Late^ models that further support our hypothesis. In the KP^Early^ model, loci enriched for Gata (lung marker), Fox (lung marker), Hnf4 (gastric marker), and Klf (pre-EMT marker) motifs were highly accessible in parental cells and less accessible after either short-term PRMT5 inhibition or in the stably resistant derivatives (**Fig. 4E–G** and **Fig. S6C**). In contrast, in the KP^Late^ model, these programs are all inactive in parental cells, display transient derepression in response to short-term PRMT5i treatment, but are then re-repressed in resistant cells (**Fig. 4G** and **Fig. S6A–C**), arguing that KP^Late^ cells briefly sample earlier developmental states but these are ultimately selected against.

We next asked whether the accessibility changes at drug-responsive loci (the bases of our HOMER analyses) persist into the stably resistant state. Our analyses showed that a high fraction of these changes (73% in KP^Early^ and 68% in KP^Late^) were retained in resistant cells (**Fig. S7D**). However, when we compare the drug-responsive loci to all resistant loci, we find that the short-term changes represent a much smaller fraction of the stable changes (21% for KP^Early^ and 11% for KP^Late^; **Fig. S7D**), indicating that resistant cells undergo substantial additional chromatin changes beyond those detected at the short-term timepoint. To characterize these additional changes, we selected loci that were significantly altered in the stable resistant state but not by short-term drug treatment, for both KP^Early^ (n=20,828) and KP^Late^ (n=25,815), divided these into repressed or derepressed relative to parental, and identified the enriched motifs using HOMER (**Fig. S7E–F**). Interestingly, these were also largely members of the previously identified KP-mutant LUAD programs (**Fig. S7E–F**). Taken together, our chromatin analyses support a model in which PRMT5 inhibition broadly expands the accessible chromatin landscape, enabling transient sampling of multiple lineage programs, followed by selection and stabilization of a subset of induced states that somehow enable drug resistance. These selected chromatin configurations converge on more dedifferentiated, aggressive disease states along the normal KP-mutant tumor progression continuum.

### A PRMT5i-induced histone methylation switch promotes chromatin accessibility and plasticity

Numerous studies report that PRMT5 mediates the symmetric dimethylation of histones^3,4,10,71–75^ and contributes to gene repression in early development^3,75^. Both H2AR3 and H4R3 have been identified as *bona fide* PRMT5 targets *in vivo*^3,4,75^. We wanted to determine whether the impact of PRMT5i on chromatin compaction in our lung cancer cell lines reflects changes in histone methylation. To address this, we isolated histones from KP^Late^-Parental cells treated with vehicle or 100 nM PRMT5i at the same 5-day timepoint used for our chromatin analyses above and conducted western blotting with an anti-H2AR3me2*s* antibody (**Fig. 5A**). This yielded a strong signal in vehicle-treated cells that colocalized with total H2A and was completely lost in the PRMT5i treated cells (**Fig. 5A**). Due to histone sequence conservation, the anti-H2AR3me2*s* antibody recognizes both H2AR3me2*s* and H4R3me2*s* (**Fig. S8A**). Thus, to further address the apparent specificity for H2AR3 in our system, we excised the histone H2A and H4 bands from SDS-PAGE gels of untreated KP^Late^-Parental histone extracts and performed liquid chromatography-tandem mass spectrometry (LC-MS/MS) analyses (**Fig. S8B–E**). We identified multiple peptides mapping to H2A and H4 (**Fig. S8B, D**), including the N-terminal ends, which displayed loss of initiator methionine residues and acetylation of the exposed N-terminus as previously reported^76,77^ (**Fig. 5B** and **Fig. S8B–E**). Importantly, a mass shift corresponding to dimethylation of R3 was detected on H2A but not H4 (**Fig. 5B** and **Fig. S8B–E**). Collectively, these data show that H2AR3 carries the H2AR3me2*s* mark in KP^Late^-Parental cells, and formation of this mark is blocked by short-term PRMT5i treatment, coincident with the reduction in chromatin compaction.

**Figure 5.**
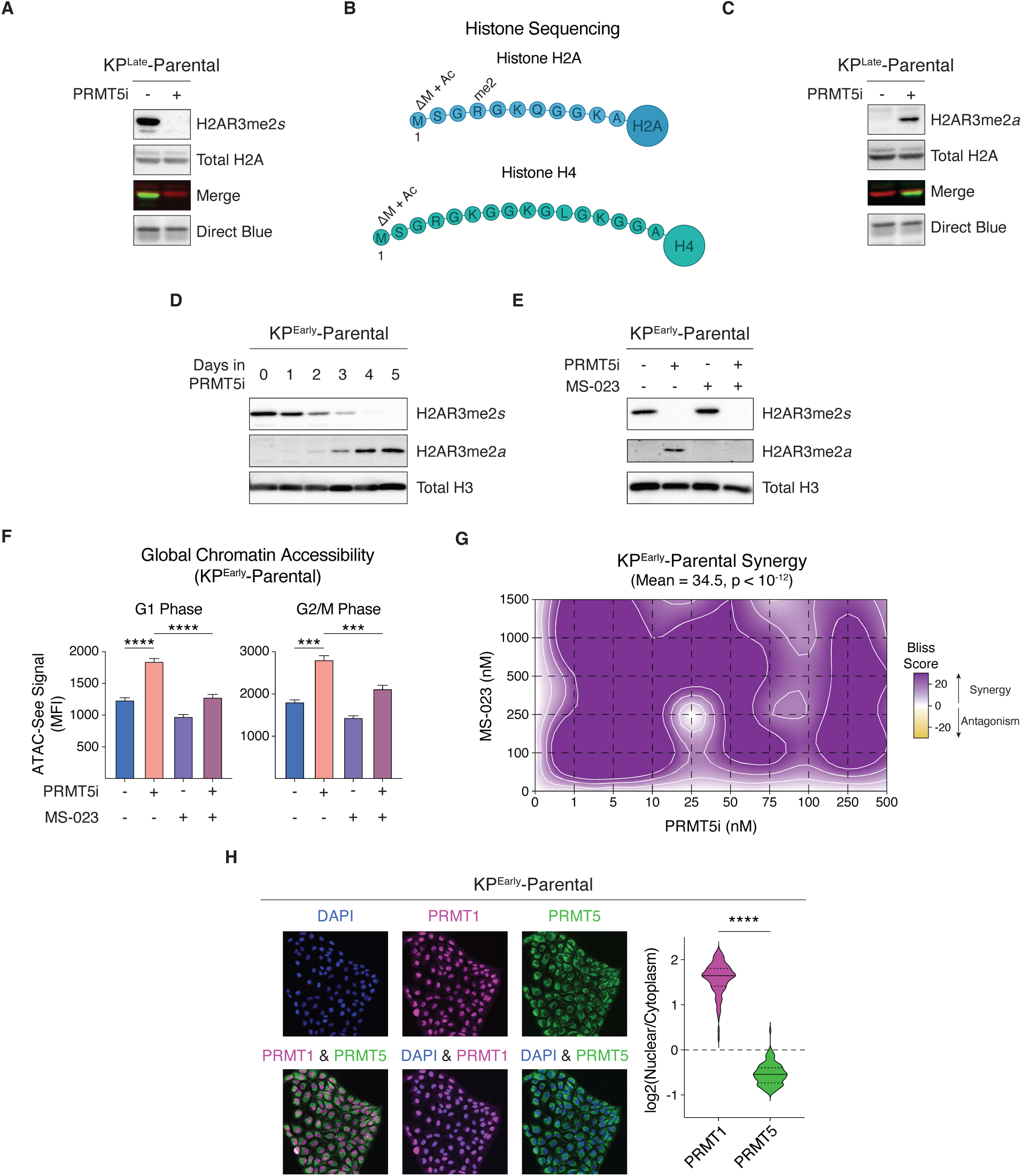
PRMT5 inhibition induces a symmetric-to-asymmetric H2AR3 methylation switch that enables chromatin plasticity. **A.** Western blot of H2AR3me2*s*, total H2A, and Direct Blue loading control in KP^Late^-Parental cells after a 5-day vehicle or PRMT5i (100 nM JNJ-64619178) treatment. The merged image shows overlap of H2AR3me2*s* (green) and total H2A (red). **B.** Schematics of N-terminal regions of H2A and H4 from KP^Late^-Parental cells sequenced by LC-MS/MS. Identified post-translational modifications are indicated above the modified residue. ΔM+Ac denotes initiator methionine loss and acetylation of the subsequent amino group, and me2 denotes dimethylation. **C.** Western blot of H2AR3me2*a*, total H2A, and Direct Blue loading control in KP^Late^-Parental cells after a 5-day vehicle or PRMT5i treatment. The merged image shows overlap of H2AR3me2*a* (green) and total H2A (red). **D.** Western blot of H2AR3me2*s* and H2AR3me2*a* in KP^Early^-Parental cells during a 5-day time course of PRMT5i treatment. Total H3 serves as a loading control. **E.** Western blot of H2AR3me2*s* and H2AR3me2*a* in KP^Early^-Parental cells after a 5-day vehicle, PRMT5i, MS-023 (type I PRMT inhibitor), or combination treatment. Total H3 serves as a loading control. **F.** Mean fluorescence intensity (MFI) of ATAC-See signal in KP^Early^-Parental cells treated with vehicle, PRMT5i, MS-023, or drug combination for 5 days. G1 and G2/M populations were assessed separately. Data are mean ± SD of 3 biological replicates per condition. **G.** Bliss synergy contour map for 5-day co-treatment of KP^Early^-Parental cells with various PRMT5i (x-axis) and MS-023 (y-axis) concentrations. Color indicates the Bliss synergy score, with yellow indicating antagonism and purple indicating synergy. **H.** Subcellular localization of PRMT1 and PRMT5 in KP^Early^-Parental cells. (*Left*) Immunofluorescence images of endogenous PRMT1 (magenta), PRMT5 (green), and DAPI (blue), with merged views indicating nuclear enrichment of PRMT1 and cytoplasmic enrichment of PRMT5. (*Right*) Violin plots depict single-cell log_2_(nuclear/cytoplasmic) intensity ratios (n=97 cells). For corresponding panels, Welch’s t-test was used. *** indicates p<0.001 and **** indicates p<0.0001.

Prior studies suggest that type I PRMTs (which catalyze ADMA formation) and type II PRMTs (which catalyze SDMA formation) can compete for shared arginine substrates, such that perturbation of one methylation type can influence the other^78–81^. As no antibody specific to H2AR3me2*a* was commercially available, we used an antibody raised against the analogous H4R3me2*a* mark (Active Motif, 39705), which we predicted would cross-react with H2AR3me2*a* based on the H2A/H4 N-terminal sequence conservation already established for the symmetric antibody (**Fig. S8A**). Western blotting with this antibody showed that H2AR3me2*a* was undetectable in vehicle-treated KP^Late^-Parental cells but robustly induced following PRMT5 inhibition (**Fig. 5C**). A similar switch from symmetric to asymmetric dimethylation of H2AR3 was observed in KP^Early^-Parental cells (**Fig. S9A**), indicating that this response is conserved across our KP-mutant LUAD models. To define the kinetics of this transition, we treated KP^Early^- and KP^Late^-Parental cells with PRMT5i for 5 days and collected histones at daily intervals. H2AR3me2*s* levels declined progressively during PRMT5i treatment and were accompanied by a reciprocal increase in H2AR3me2*a* (**Fig. 5D** and **Fig. S9B**), consistent with a model in which the loss of PRMT5-dependent methylation permits subsequent asymmetric dimethylation of H2AR3 by type I PRMTs. To test this directly, we treated cells with the pan-type I PRMT inhibitor MS-023, either alone or in combination with PRMT5i. MS-023 alone had no effect on H2AR3me2*s* levels but completely abrogated the accumulation of H2AR3me2*a* induced by PRMT5 inhibition (**Fig. 5E** and **Fig. S9C**), demonstrating that the methylation switch is dependent on type I PRMT activity.

Because histone ADMA has been broadly associated with chromatin opening and transcriptional activation^82–86^, whereas SDMA is linked to transcriptional repression^3,4^, we asked whether this methylation switch contributes to PRMT5i-induced chromatin remodeling. KP^Early^- and KP^Late^-Parental cells were treated with vehicle, PRMT5i, MS-023, or the combination for 5 days and subjected to ATAC-See analysis. Consistent with our prior results, PRMT5i induced widespread chromatin opening in both cell lines (p<0.001; **Fig. 5F** and **Fig. S9D**). In contrast, MS-023 reduced chromatin accessibility when administered alone, and it significantly blunted the accessibility gains induced by PRMT5 inhibition (p<0.01; **Fig. 5F** and **Fig. S9D**). These findings suggest that the PRMT5i-induced switch from the symmetric (H2AR3me2*s*) to the asymmetric (H2AR3me2*a*) mark on H2AR3 contributes to global chromatin remodeling. We then asked whether type I PRMT activity was actively contributing to resistance acquisition. To address this, we compared the viability of KP^Early^- and KP^Late^-Parental cells in response to a range of PRMT5i and/or MS-023 concentrations (**Fig. 5G** and **Fig. S9E–F**). This revealed strong synergy between PRMT5i and MS-023, with mean Bliss scores of 34.5 and 22.7 for KP^Early^- and KP^Late^-Parental cells, respectively. Collectively, these data establish type I PRMT activity as a functional driver of the adaptive chromatin response and a potential co-therapeutic target to limit resistance.

Given these findings, we then asked how H2AR3me2*s* predominates in untreated cells and is replaced by H2AR3me2*a* following PRMT5 inhibition. RNA-Seq analyses revealed that PRMT5 and PRMT1, the major type I PRMT, are both highly expressed in KP^Early^- and KP^Late^-Parental cells and their levels are unchanged by PRMT5i treatment (**Fig. S9G**), arguing against a transcriptional basis for the switch. We then examined the subcellular localization of these enzymes by immunofluorescence. In both cell lines, a substantial fraction of PRMT5 localized to the cytoplasm, whereas PRMT1 was almost exclusively nuclear (p<0.0001; **Fig. 5H** and **Fig. S9H– I**). This spatial segregation provides a simple explanation for both the dominance of H2AR3me2*s* in untreated cells and the rapid emergence of H2AR3me2*a* following PRMT5 inhibition. Under basal conditions, PRMT5 can methylate newly synthesized histones in the cytoplasm before their nuclear import and exposure to PRMT1. Upon PRMT5 inhibition, newly synthesized histones enter the nucleus with unmodified H2AR3 and become substrates for PRMT1-mediated modification, enabling a widespread switch from H2AR3me2*s* to H2AR3me2*a* that replaces chromatin repression with activation to drive cellular plasticity.

### *MTAP* loss engages the histone methylation switch, and *CDKN2A/MTAP*-deficient human tumors are more dedifferentiated

The clinical utility of early-generation PRMT5i is limited by on-target toxicities in non-cancerous proliferative tissues^87^. Newer-generation PRMT5i were developed to overcome this by targeting cancer cells that harbor loss of *MTAP* (methylthioadenosine phosphorylase) due to its adjacency to the frequently deleted *CDKN2A* tumor suppressor locus (**Fig. 6A–B**). MTAP functions in the SAM-generation cycle and its loss causes accumulation of its substrate, methylthioadenosine (MTA), which inhibits PRMT5^88,89^ (**Fig. 6A**). These MTA-cooperative inhibitors form a PRMT5i•MTA•PRMT5 ternary complex that selectively inhibits PRMT5 activity in *CDKN2A*/*MTAP*-deficient tumors^21–23,90^.

**Figure 6.**
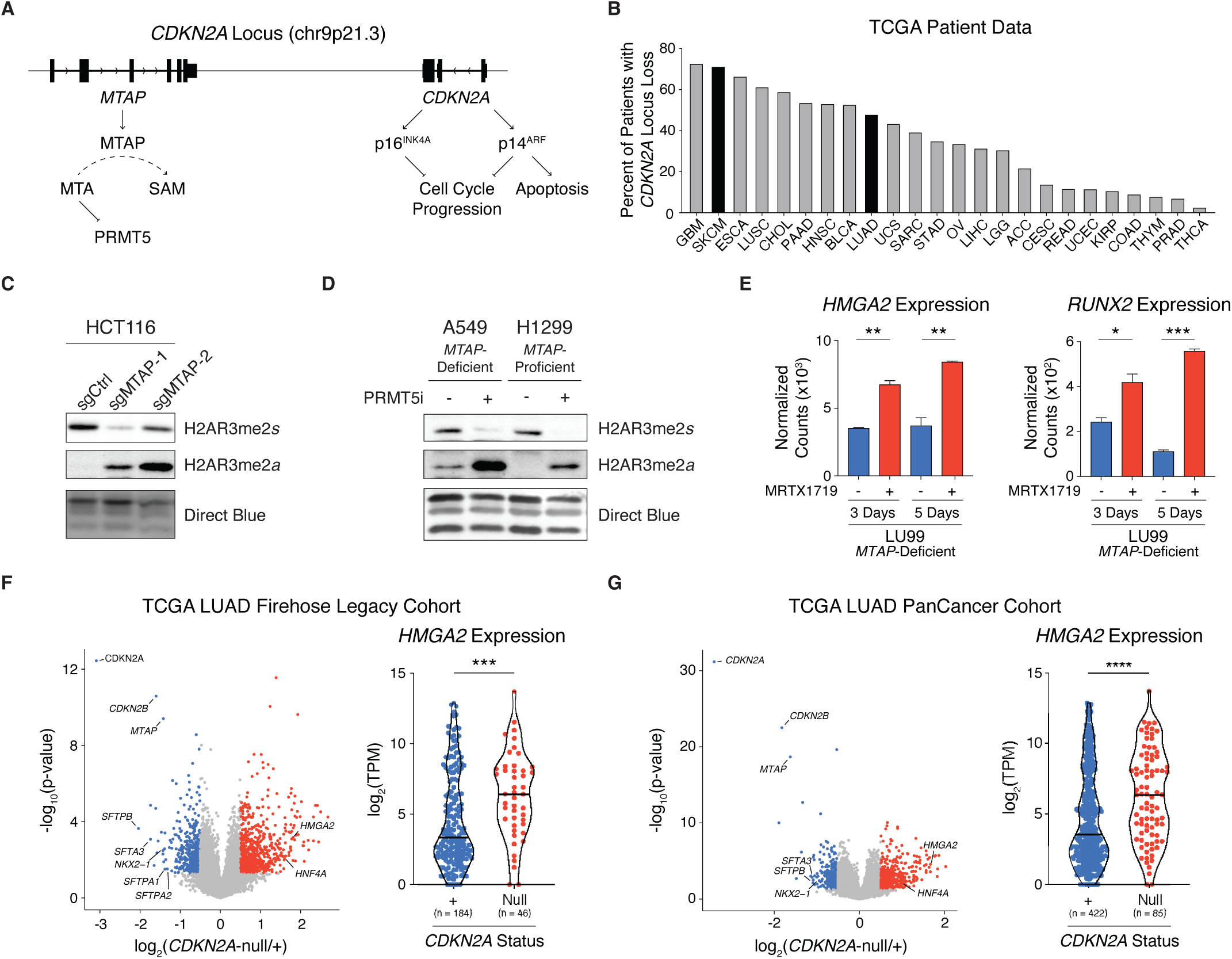
MTAP loss engages the histone methylation switch and CDKN2A/MTAP-deficient tumors are more dedifferentiated. **A.** Schematic of the *CDKN2A* and adjacent *MTAP* loci indicating gene products and functions. **B.** Frequencies of *CDKN2A* copy number losses in various tumor types, with SKCM (skin cutaneous melanoma) and LUAD highlighted. **C.** Western blot of H2AR3me2*s*, H2AR3me2*a*, and Direct Blue loading control in control HCT116 cells and two independent CRISPR-generated *MTAP*-knockout clones. **D.** Western blot for H2AR3me2*s*, H2AR3me2*a*, and Direct Blue loading control in *CDKN2A*/*MTAP*-deficient A549 and *MTAP*-proficient H1299 human LUAD cells treated with vehicle or 2.5 nM PRMT5i for 5 days. **E.** Normalized RNA expression counts for *HMGA2* and *RUNX2* in *CDKN2A/MTAP*-deficient LU99 cells treated with vehicle (blue) or MRTX1719 (red), an MTA-cooperative PRMT5i, for 3 or 5 days. Data are mean ± SD of 2 biological replicates per condition. **F–G.** Differentially expressed genes comparing *CDKN2A*-null to the rest of the tumors (*CDKN2A*+) in the TCGA LUAD (**F**) Firehose Legacy or (**G**) PanCancer cohorts. (*Left*) Points represent individual genes that have increased expression (red, p<0.05, log_2_FC>0.5) or decreased expression (blue, p<0.05, log_2_FC<–0.5). (*Right*) Expression of *HMGA2* in indicated groups. Points represent individual patients with the total number of patients per group listed below the axis. For corresponding panels, Welch’s t-test was used. * indicates p<0.05, ** indicates p<0.01, *** indicates p<0.001, and **** indicates p<0.0001.

We wondered whether *MTAP* loss is sufficient to engage the histone methylation switch identified above in the basal state. To test this, we took advantage of two independent, isogenic HCT116 *MTAP*-knockout clones we previously generated^12^. Western blotting revealed a decrease in H2AR3me2*s* and a corresponding increase in H2AR3me2*a* in both of the *MTAP*-knockout clones, compared to their parental control (**Fig. 6C**). Notably, consistent with the fact that MTA accumulation yields partial suppression of PRMT5 activity, this effect is more subtle than the near-complete switch from H2AR3me2*s* to H2AR3me2*a* that results from pharmacological PRMT5 inhibition (**Fig. 5A** and **C**). Thus, *MTAP* loss alone is sufficient to partially engage the same histone methylation switch that drives chromatin plasticity in our KP-mutant LUAD model, and it does so in colorectal cancer cells, a context entirely distinct from lung adenocarcinoma.

We next asked whether the methylation switch, and the impact of *MTAP* status, extend to human LUAD, by examining H2AR3me2*s* and H2AR3me2*a* in A549 cells (which are *CDKN2A*/*MTAP*-deficient) and H1299 cells (*MTAP*-proficient) after a 5-day treatment with vehicle or 2.5 nM PRMT5i (**Fig. 6D**). In the absence of drug, we detected a baseline H2AR3me2*a* signal specifically in the *MTAP*-deficient A549 line, and not the H1299 cells, and PRMT5i treatment yielded a near-complete switch from H2AR3me2*s* to H2AR3me2*a* in both contexts. Collectively, these data argue that *MTAP* loss shifts the balance of histone methylation toward asymmetric modification prior to any pharmacological intervention, and complete PRMT5 inhibition drives the switch fully irrespective of *MTAP* status.

Having shown that *MTAP* loss primes the histone methylation switch, we asked whether MTA-cooperative inhibitors, which specifically suppress residual PRMT5 activity in this genetic background, serve to induce LUAD dedifferentiation phenotypes. For this, we used a published RNA-Seq dataset^23^ from *CDKN2A*/*MTAP*-deficient LU99 cells treated for 3 or 5 days with MRTX1719, an MTA-cooperative PRMT5i in clinical trials. We found that core late-stage disease markers *HMGA2* and *RUNX2* were both significantly upregulated by 3-day treatment (p<0.01 and p<0.05, respectively) and further elevated by 5-day treatment (p<0.01 and p<0.001, respectively; **Fig. 6E**). Thus, MTA-cooperative PRMT5i also trigger the dedifferentiation phenotype we uncovered with early-generation PRMT5i.

In analyzing these data, we noted that the magnitude of dedifferentiation marker induction in response to short-term treatment in LU99 cells (**Fig. 6E**) was more profound than the short-term responses of our KP^Early^ and KP^Late^ lines. We speculated that this could reflect partial engagement of the symmetric-to-asymmetric methylation shift in the basal, untreated state of the *MTAP*-deficient LU99 cells. Considering this in a broader context, we wondered whether *CDKN2A/MTAP*-deficient tumors tend to be more dedifferentiated than their proficient counterparts. We used TCGA patient data to address this question. We began with two different LUAD patient cohorts, identifying tumors that harbored homozygous *CDKN2A* deletion (*CDKN2A*-null) and comparing these with the rest of the cohort (*CDKN2A*+). We confirmed that *MTAP* expression was significantly reduced in the *CDKN2A*-null tumors, and then examined LUAD differentiation markers (**Fig. 6F–G** and **Fig. S10A–B**). Consistent with our hypothesis, the *CDKN2A*-null tumors were significantly more dedifferentiated than their *CDKN2A*+ counterparts, as they expressed lower levels of *NKX2-1* (p<0.05) and alveolar surfactant proteins (including *SFTPB* [p<0.01]) and higher levels of *HMGA2* (p<0.001).

Importantly, *CDKN2A/MTAP* deficiency, as well as the efficacy of PRMT5i, is seen across a broad spectrum of tumor types. We therefore extended our analyses to a second cancer type: cutaneous melanoma (SKCM), in which *CDKN2A* loss is especially prevalent (**Fig. 6B**) and disease progression is marked by a transition from melanocyte identity in early-stage tumors to undifferentiated, neural-crest-like states in advanced disease^91^. We examined two different melanoma TCGA datasets, again confirming that *MTAP* expression was significantly reduced in *CDKN2A*-null versus *CDKN2A*+ tumors (**Fig. S10C–D**). We then examined differentiation markers, and found significantly lower levels of melanocyte markers, including melanin synthesis genes, such as *OCA2*^91^ (p<0.001), and higher levels of undifferentiated and neural crest markers, such as *EDN3*^92^ (p<0.001), in the *CDKN2A*-null melanomas (**Fig. S10C–D**). Taken together, our data demonstrate that *CDKN2A*/*MTAP* deficiency is associated with a higher degree of dedifferentiation in human tumors.

## Discussion

We have previously shown that KP-mutant LUAD cells develop rapid resistance to PRMT5i through an adaptive response rather than selection of pre-existing resistant variants^28^. Our current study establishes the molecular events that drive acquisition of that resistance and examines the consequences of that disease state. Unexpectedly, we find that resistance does not mitigate the splicing defects that are responsible for cancer cell vulnerability to PRMT5i^5,12,13^. Specifically, PRMT5i treatment causes resistant variants to display the same splicing changes as their parental controls, with regard to both identity and magnitude of change, despite their profoundly improved survival. Instead, resistance reflects a developmental state shift that, in the case of our KP-mutant LUAD model, corresponds to a drastic shift along the normal KP-mutant tumor progression continuum toward metastatic states. Importantly, PRMT5i actively enables this state shift through its influence on histone regulation, driving a switch from the transcriptionally repressive H2AR3me2*s* mark to the transcriptionally active H2AR3me2*a* mark, thereby inducing widespread chromatin opening. The resulting derepression of lineage programs enables sampling and selection of the state(s) best suited to cell survival. Notably, the link between PRMT5 inhibition and dedifferentiation extends beyond our mouse model: a human LUAD cell line shows the same dedifferentiation response to a clinical-stage, tumor-selective MTA-cooperative PRMT5i, and human *CDKN2A*/*MTAP*-deficient LUAD and melanoma tumors, which carry endogenous PRMT5 suppression, display a correspondingly higher degree of dedifferentiation. In essence, PRMT5 inhibition simultaneously creates a splicing-dependent vulnerability and activates a chromatin-mediated adaptive program – two inseparable consequences of targeting PRMT5. This coupling imposes a fundamental constraint on therapeutic efficacy: the same intervention that induces lethality also launches the mechanism of escape.

Our data are consistent with a model in which the chromatin changes induced by PRMT5i result from a symmetric-to-asymmetric methylation switch on H2AR3, in which loss of cytoplasmic PRMT5 activity allows previously inaccessible substrate to become available to nuclear PRMT1. Our model is the converse of a reciprocal relationship reported previously, in which loss of PRMT1 unmasks substrates for scavenging by type II PRMTs^78^. That prior work, however, did not examine histones and concluded PRMT1 is simply the dominant enzyme whose pre-existing mark masks substrates from other methyltransferases – a passive, abundance-driven process. Our finding that PRMT5i unleashes PRMT1 activity on histones argues that this competition is not passive masking but hierarchical, governed by ordered, compartmentalized access to the substrate – with PRMT5, the dominant enzyme at this site, encountering histones first. Newly synthesized histones are initially released into the cytoplasm, where PRMT5 resides, giving it the first opportunity to modify H2AR3. Histones only encounter PRMT1 after nuclear entry, ensuring that ADMA modification can only occur if SDMA was not written or was actively removed. Consistent with this model, we find that our untreated cells have high levels of H2AR3me2*s* and almost no H2AR3me2*a*, even though these cells express significant levels of PRMT1. This ensures continued chromatin repression and silencing of any lineage-specification programs that are inappropriate for the basal cell state. Interestingly, PRMT5 subcellular localization is known to be developmentally regulated: this enzyme actively shifts between cytoplasmic and nuclear compartments during early embryogenesis^3^. This strongly suggests that the hierarchy between PRMT5 and PRMT1 is used as a tunable function at specific developmental stages, presumably to control lineage choices.

In our cells, PRMT5 inhibition results in a rapid loss of H2AR3me2*s* and a reciprocal rise in H2AR3me2*a*, which promotes widespread chromatin remodeling. Since co-treatment with MS-023 almost completely abolishes this chromatin opening, we can conclude that SDMA loss alone is insufficient to drive remodeling and that active engagement of type I PRMTs, likely PRMT1, is required. The consequent widespread gene re-activation offers the perfect opportunity for selection of drug resistance. In the case of targeted cancer therapies, resistance typically results from selection of rare, pre-existing cells whose molecular features confer drug tolerance. The mechanism for PRMT5i is fundamentally different. Rather than selecting pre-existing resistant cells, the drug actively generates new epigenetic diversity – presumably a range of accessible chromatin configurations – and selection then acts on that diversity to yield resistant states. After short-term drug treatment, the enriched TF motifs in differentially accessible regions lie on the normal KP-mutant LUAD progression continuum. It is unclear whether the induced changes are restricted to this chromatin landscape, or whether other lineage specification programs are activated but simply miss the threshold for significance. Regardless, KP-mutant LUAD programs dominate the accessibility states available for selection. In KP^Early^ cells, many drug-induced changes were maintained in the resistant state, consistent with the parental cells starting at an earlier stage of the differentiation continuum and sampling, and ultimately selecting, more advanced, dedifferentiated cell states. The behavior of KP^Late^ cells is more complex, and provides the strongest support for the selection hypothesis. These cells start at a later stage of the KP-mutant LUAD continuum, and short-term PRMT5i treatment transiently activates many programs, including markers of earlier disease states. However, these transiently activated early-stage programs are subsequently repressed in the stably resistant state. Importantly, despite the differing starting points of KP^Early^ and KP^Late^ cells, their stably resistant variants converge on a shared transcriptional state. It is unclear how this state helps escape the splicing defects induced by PRMT5i treatment, but it follows a recurring theme in cancer biology where acquisition of more embryonic/stem cell fates enables drug tolerance to many treatments^93,94^.

Our findings have serious implications for cancer patients. The PRMT5i-resistant variants are significantly more metastatic than their parental counterparts. Moreover, we have shown that a modest two-week treatment with PRMT5i *in vivo* yields significant increases in tumor grade. We observed no reduction in lung tumor burden, whereas tumor regression has been reported in other PRMT5i studies^20,21,23^. Critically, the grade advancement we observe does not depend on any effect on tumor burden, and our data therefore indicate that PRMT5i actively promotes progression regardless of whether it debulks tumors in a given context. Given the rapidity of this grade advancement and the fact that chromatin remodeling is inextricably coupled to the splicing changes that make PRMT5i effective in the first place, this warrants careful consideration in the clinical development of PRMT5i. Essentially, the deleterious consequences of PRMT5i action – driving tumor cells toward a more aggressive and treatment-resistant state – may well outweigh any tumor-suppressive effects. We envisage two possible ways to bypass this problem. The optimal solution would be to specifically target PRMT5’s role in splicing, without disrupting its role in histone regulation. We have achieved this genetically, by deleting the co-factor essential for PRMT5’s splicing role and showing that this kills cancer cells as effectively as PRMT5 inhibition^12^. Unfortunately, there is currently no small molecule that achieves this goal. The alternative approach, as supported by our current study, would be to try dual targeting of PRMT5 and type I PRMTs, suppressing chromatin remodeling at its enzymatic root while enhancing cytotoxicity. Whether this combination is sufficient to prevent the progression phenotype without driving toxicity remains an important question for future work.

While our study focused on KP-mutant LUAD, PRMT5 inhibition is effective in many cancer contexts^5,12,13,20–23^. As long as these tumors express type I PRMTs, we expect PRMT5i treatment to trigger changes in the methylation state of H2AR3 and induce chromatin remodeling. If dedifferentiation is typically achieved, drug resistance could be a frequent outcome. Our analysis of *CDKN2A*/*MTAP*-deficient tumors provides support for this notion: these tumors carry chronically reduced PRMT5 activity, via endogenous inhibition by elevated MTA levels, and we find this is associated with a higher degree of dedifferentiation in both LUAD and cutaneous melanoma. This concern is not limited to PRMT5; many chromatin-modifying enzymes in development and/or clinical trials act through analogous mechanisms of writing, erasing, or reading histone marks that constrain lineage identity. These agents, by definition, have the potential to rewire the chromatin landscape. Whether drug-induced chromatin plasticity is a general mechanism of resistance across epigenetic therapies, and whether the resulting resistant states consistently favor more aggressive disease, are questions that warrant careful investigation.

## Materials and Methods

### Cell Line Generation

Parental, control, and drug-resistant derivatives of murine KP LUAD cell lines KP393T5 (KP^Late^) and KP1233 (KP^Early^) were previously described^28,29,95^. Cells were cultured in Dulbecco’s Modified Eagle’s Medium (DMEM) supplemented with 10% fetal bovine serum (FBS) and 1% penicillin-streptomycin at 37°C with 5% CO_2_. Cells were routinely tested for mycoplasma and were found to be negative. For luciferase expression, cells were infected with lentivirus containing pLenti Luciferase-P2A-mCerulean and sorted on mCerulean expression to generate stable cell lines. HCT116, H1299 and A549 cells were obtained from ATCC. Generation of *MTAP*-deficient HCT116 cells was previously described^12^.

### Dose-Response Curves

The PRMT5 inhibitor (PRMT5i) JNJ-64619178 was used for all treatment experiments (unless otherwise noted) and obtained from MedChemExpress (HY-101564). An additional PRMT5i, EPZ015666, was obtained from Levi Garraway. For dose response curves, 100 cells were plated in a volume of 150 μL media into each well of a 96-well plate. Perimeter wells were filled with 200 μL of media to avoid edge effects. The following day, 50 μL of 4× the final drug concentration in media was added to each well in triplicate. For synergy analyses, 100 cells were plated in a volume of 100 μL media into each well of a 96-well plate. The following day, 50 μL of 4× the final drug concentration in media was added to each well in triplicate for each inhibitor. After 5 days, plates were washed with PBS, and 50 μL 0.5% crystal violet in 20% methanol was added and allowed to incubate for 20 minutes on an orbital shaker. Plates were submerged in water and allowed to air dry. For imaging, 200 μL of 100% methanol was added to each well, and plates were incubated for 20 minutes on an orbital shaker. The optical density of each well at 570 nm (OD_570_) was measured with a Tecan M200 Pro plate reader. Background signal was determined by averaging the OD_570_ values of cell-free wells and was subtracted from all viability measurements. Average signal for each dose triplicate was computed, and drug-treated cells were normalized to their DMSO control by division. To generate curves, normalized viability values for each sample were fit to the following nonlinear model:

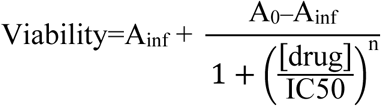

where A_inf_ and A_0_ are the lowest and highest normalized viability values, respectively. Area under the curve was computed and used for statistical testing. Bliss synergy scores were calculated from normalized viability using SynergyFinder Plus^96^.

### RNA Sequencing and Differential Gene Analysis

#### Library Preparation and Sequencing

KP^Early^- and KP^Late^-Parental and Resistant lines were treated for 3 days with either DMSO (vehicle) or PRMT5i (JNJ-64619178 for KP^Early^ and EPZ015666 for KP^Late^). Following treatment, cells were harvested by scraping. RNA was purified using the RNeasy Mini Kit (Qiagen, 74106) according to manufacturer’s instructions, including the optional DNaseI addition to remove contaminating gDNA (Qiagen, 79254).

For KP^Early^ cells, RNA-Seq libraries were prepared using the NEBNext Ultra II Directional RNA Kit with Poly(A) Selection and sequenced using the Element AVITI sequencer, with 75 nucleotide paired-end reads. For KP^Late^ cells, RNA-Seq libraries were prepared using the NEBNext Ultra II Directional RNA Kit with Poly(A) Selection and sequenced using the NextSeq 500, with 75 nucleotide paired-end reads.

#### Alternative Splicing Analysis

Raw RNA-Seq reads were mapped using STAR aligner version 2.5.3a^97^ with the following parameters:

~~~
STAR
--runMode alignReads
--runThreadN 16
--sjdbOverhang 149
--outReadsUnmapped Fastx
--outSAMtype BAM SortedByCoordinate
--outFilterMultimapNmax 20
--outFilterMismatchNmax 999
--outFilterMismatchNoverLmax 0.04
--alignIntronMin 70
--alignIntronMax 500000
--alignMatesGapMax 500000
--alignSJoverhangMin 8
--alignSJDBoverhangMin 1
--outSAMstrandField intronMotif
--outFilterType BySJout
--twopassMode Basic
~~~

Additionally, raw RNA-Seq reads were mapped using Bowtie 1.2.3^98^ using the following parameters:

~~~
bowtie -n 1 -l 30 -e 70 -t -p 4 -m 1 -X 500 -S
~~~

Detained intron and alternative splicing event identification was performed as previously described^5,14^, with the following modifications: Mapped reads were filtered using Bedtools v2.26.0 to remove reads overlapping expressed repeats, annotated in the UCSC Genome Browser RNA repeats track and Gencode mm10, as well as coding exons annotated in Gencode mm10. Alternative and constitutive classifications were performed using custom Python scripts and are agnostic with regard to existing annotations other than known gene boundaries. Importantly, all cell lines were run in parallel to ensure that the annotations would be consistent between the cell lines.

#### Detained Intron and Alternative Splicing Event Quantification (DEXSeq)

Detained intron and alternative splicing events were quantified using DEXSeq^99^. A genome annotation including all alternative splicing events and detained introns was generated using custom Python scripts and Python scripts that accompany DEXSeq. Reads were then quantified to this genome using the DEXSeq Python script with the following parameters:

~~~
-p yes -s reverse -f bam -r pos
~~~

Finally, differential splicing was quantified using the DEXSeq package within R. Alternative splicing events and detained introns were called as significant if p_adj_<0.05 and lowly expressed exons and introns were filtered out if their normalized counts were below 80.

#### Detained Intron and Coding Isoform Quantification (RSEM-DESeq2)

Transcript annotation was performed as previously described^5^. Briefly, using custom Python scripts, we derived a complete transcriptome annotation based on Gencode mm10 gene start and end boundaries and the location of poly(A) sites, the genomic locations of detained introns, and the genomic locations of all splice junctions (from the STAR alignment). For comparisons of isoforms containing detained introns to the consensus coding sequence, the annotation for each gene was set to identify the consensus isoform, i.e., the junctions defining each exon are the most frequently used junction detected within all of the combined samples. Each detained intron within a gene was then assigned to an additional transcript.

#### Gene and Isoform Expression

Isoform-level quantification was performed using RSEM v1.3.1^39^ using the custom genomic annotation described above, as well as Gencode mm10 annotations with the parameters:

~~~
rsem-calculate-expression --forward-prob 0 --no-bam-output --paired-end
~~~

Differential expression analysis of gene and isoform count tables was performed using DESeq2 v1.42.0^100^. Normalized count tables were obtained from DESeq2. Gene and isoform expression differences were considered significant if p_adj_<0.05.

#### Gene Ontology Analysis

Gene Ontology (GO) analysis was conducted using DAVID^37,38^. Genes that contained a PRMT5i-responsive DI (p_adj_<0.05) were run against the KW Biological Process category. A hypergeometric test was used to determine the probability of observing a certain overlap between the inputted gene list and a specific GO term, compared to the background gene list. For these comparisons, the background gene list was all genes in the mouse genome. To account for multiple hypothesis testing, we report the Benjamini-Hochberg-corrected FDR. Gene sets were considered significant if the corrected FDR<0.01. Gene sets are displayed if they met the significance threshold in at least one cell line.

#### Differential Gene Expression Analysis

Reads were aligned and quantified using the NF-Core RNA-Seq pipeline v3.12.0^101^, using the following optional flags:

~~~
--gencode
--aligner star_salmon
--pseudo_aligner salmon
~~~

Differential gene expression was run on the STAR-Salmon count tables using DESeq2^100^, and normalized RNA counts were obtained using the `SizeFactor` generated by DESeq2. PCA was run using the 500 most variable genes. Pre-ranked gene set enrichment analysis was performed using the test statistic as the ranking metric, and samples were run against the C2 and C5 MSigDB collections^40^.

### Quantitative Real-Time PCR (qRT-PCR)

Total RNA was purified from cultured cells using the RNeasy Mini Kit (Qiagen, 74106) according to manufacturer’s instructions, including the optional DNaseI addition (Qiagen, 79254). RNA was converted to cDNA using SuperScriptIII (Thermo Fisher Scientific, 18080044). Quantitative PCR reactions were performed using FAST-SYBR Green (Thermo Fisher Scientific, 4385612) on a StepOnePlus Real-Time PCR System (Applied Biosystems). Data were analyzed using the ΔΔC_T_ method, and relative messenger RNA (mRNA) levels were normalized to total gene levels. The following primers were used:

*Sf3b2* Detained Intron F: AGCTCTGAAATGGACTCTAAC
*Sf3b2* Detained Intron R: GCTACACGATGCCTTCTT
*Sf3b2* Total Gene F: CTTGTGCTCCTCCTCAAA
*Sf3b2* Total Gene R: GGCTTTCAAGCTAACTGATG
*Sos2* Detained Intron F: TCAATAGTGGAAGGGTTTGA
*Sos2* Detained Intron R: ATCAGCGCAGATCTGAAA
*Sos2* Total Gene F: TCATGGCTTTCCCTCTTAC
*Sos2* Total Gene R: ACTGGGTTGAGCATCATT

### Western Blot

Cells were harvested by scraping, washed with PBS, and lysed in RIPA buffer (50 mM Tris pH 8.0, 150 mM NaCl, 1% NP-40, 0.5% sodium deoxycholate, and 0.1% sodium dodecyl sulfate) supplemented with EDTA-free protease inhibitor cocktail (MilliporeSigma, 11836170001). For drug treatment experiments, cells were treated with DMSO (vehicle) or 100 nM PRMT5i (JNJ-64619178) for 5 days, before harvesting, unless otherwise indicated.

Protein concentration was determined using the Pierce BCA Protein Assay Kit (Thermo Fisher Scientific, 23227), then 30 μg of protein was mixed with 4× Laemmli buffer and loaded onto 12% SDS-polyacrylamide gels. Proteins were separated by SDS-PAGE, transferred onto nitrocellulose membranes, and probed overnight at 4°C in 5% non-fat milk in TBST (unless otherwise stated) with antibodies against each of the following: NKX2-1 (Abcam, ab76013, 1:1,000), HNF4A (Cell Signaling Technology, 3113S, 1:1,000), HMGA2 (Thermo Fisher Scientific, PA5-21320, 1:1,000), SDMA (Cell Signaling Technology, 13222S, 1:1,000, 5% BSA), HSP90 (BD, 610418, 1:2,000), GAPDH (Thermo Fisher Scientific, AM4300, 1:5,000). Membranes were then incubated for 1 hour at room temperature with anti-mouse (LI-COR Biosciences, 926-32210, 1:10,000) or anti-rabbit (LI-COR Biosciences, 926-32211, 1:10,000) conjugated to IRDye 800CW and fluorescence was measured using the ChemiDoc Imaging System (Bio-Rad).

### Histone Extraction

Unless otherwise indicated, cells were treated with DMSO (vehicle) or 100 nM PRMT5i (JNJ-64619178) for 5 days. Exceptions were as follows: A549 and H1299 cells were treated with 2.5 nM JNJ-64619178; for time-course experiments, cells were treated with 100 nM JNJ-64619178 and histones collected at 24-hour intervals; for type I PRMT inhibition experiments, cells were treated with 100 nM JNJ-64619178, 1 μM MS-023, or the combination for 5 days; and HCT116 parental and *MTAP*-knockout cells were grown without drug treatment. Cells were harvested in cold PBS. Packed cell volume (typically ∼50 μL) was used to scale all subsequent steps. Cells were washed twice in Nuclear Isolation Buffer (NIB; 15 mM Tris-HCl pH 7.5, 15 mM NaCl, 60 mM KCl, 5 mM MgCl_2_, 1 mM CaCl_2_, 250 mM sucrose) supplemented fresh with protease inhibitors, followed by centrifugation at 1,000 × *g* for 5 min at 4°C. Cell pellets were lysed in NIB containing 0.3% NP-40 alternative (10× pellet volume), vortexed, and incubated on ice for 5 min to isolate nuclei. Nuclei were pelleted (1,000 × *g*, 5 min, 4°C), washed twice with NIB lacking detergent, and resuspended in 0.2 M H_2_SO_4_ (5× pellet volume). Acid extraction was performed for 2–4 hours at 4°C with rotation, followed by centrifugation at 3,400 × *g* for 5 min at 4°C to collect the supernatant. Histones were precipitated by addition of trichloroacetic acid (TCA; final ∼20– 30%) and incubated overnight at 4°C. Precipitated proteins were pelleted (3,400 × *g*, 5 min, 4°C), washed sequentially with ice-cold acetone containing 0.1% HCl and then acetone alone, and briefly air-dried. Pellets were resuspended in 50 mM ammonium bicarbonate (30–50 μL).

Protein concentration was determined using the Pierce BCA Protein Assay Kit (Thermo Fisher Scientific, 23227), then 6 μg of protein was mixed with 4× Laemmli buffer and loaded onto SDS-polyacrylamide gels. For analysis of dimethylated recombinant nucleosomes, 2 μg of nucleosomes containing either H2AR3me2*s* (Epicypher, 16-0361) or H4R3me2s (Epicypher, 16-0358) were loaded onto SDS-polyacrylamide gels. Proteins were separated by SDS-PAGE and transferred onto nitrocellulose membranes. Following transfer, total histone loading was visualized by staining membranes in 0.008% (w/v) Direct Blue 71 (MilliporeSigma, 212407) in 40% ethanol / 10% acetic acid for 5 minutes and rinsing in TBST prior to blocking. Membranes were then probed overnight at 4°C in 5% non-fat milk in TBST (unless otherwise stated) with antibodies against each of the following: H2AR3me2*s* (PTM BIO, PTM-673, 1:1,000), H4R3me2a (Active Motif, 39705, 1:1,000), Histone H2A (Cell Signaling Technology, 3636S, 1:1,000), Histone H3 (Cell Signaling Technology, 4499S, 1:1,000). Membranes were then incubated for 1 hour at room temperature with anti-mouse (LI-COR Biosciences, 926-68070, 1:10,000) conjugated to IRDye 680RD and/or anti-rabbit (LI-COR Biosciences, 926-32211, 1:10,000) conjugated to IRDye 800CW and fluorescence was measured using the ChemiDoc Imaging System (Bio-Rad).

### scRNA-Seq Analysis

Processed scRNA-Seq data^32^ were downloaded from the Broad Institute Single Cell Portal. Plots were rendered using Seurat v5.0.1.

### ATAC-Sequencing

Cells were treated with DMSO (vehicle) or 100 nM PRMT5i (JNJ-64619178) in biological triplicate, with independently plated cultures harvested and processed separately for library preparation. After 5 days, 1×10^5^ cells were harvested for nuclear isolation, transposition, and library preparation as previously described^102^. The pooled Illumina library was converted to Element AVITI chemistry. Libraries were sequenced using the Element AVITI sequencer, with 75 nucleotide paired-end reads. Raw reads were aligned and quantified using the NF-Core ATAC-Seq pipeline v2.1.2^101^, using the following optional flags:

~~~
--genome mm10
--blacklist ENCFF547MET.bed
--min_reps_consensus 3
--mito_name chrM
--macs_gsize 2652783500
--narrow_peak
~~~

Differential accessibility was run on the MACS count tables using DESeq2, and normalized peak counts were obtained using the `SizeFactor` generated by DESeq2. Normalized BigWig accessibility tracks were visualized using IGV v2.8.2. PCA was run using the 5,000 most variable peaks. Peaks were called differentially accessible if they met the following criteria: p_adj_<10^−4^, |log_2_FC|>0.5 for parental/resistant analysis, or p_adj_<0.01, |log_2_FC|>1 for drug treatment analysis.

TF motif analysis was performed using chromVAR^54^. In brief, 300 bp peak windows were compared to annotated mouse TF motifs obtained from the “mouse_pwms_v2” database to generate a binary overlap matrix. Background peaks were sampled (n=500 iterations) to adjust for GC bias. Motif accessibility was computed using the `computeDeviations` function. For TF family deviation scores, individual TF motifs with 75% or greater correlation were grouped for deviation and variance analyses.

Accessibility heatmaps were generated using deepTools v3.0.1^103^. BED files were generated for each peak group. Peak matrices were generated for each sample and peak group using `computeMatrix` with the optional flags:

~~~
--referencePoint center
-b 500
-a 500
-bs=1
-p=max
--missingDataAsZero
~~~

ATAC-Seq heatmaps were generated using plotHeatmap with the optional flags:

~~~
--zMin 0.3
--zMax 1.5
--missingDataColor 1
~~~

### TF Motif Enrichment Analysis

BED files for each peak set were generated. *De novo* motif discovery was performed using HOMER v4.10 (`findMotifsGenome.pl`)^67^ using a 300 bp window, with enrichment calculated against HOMER’s default GC-matched genomic background. *De novo* motifs were considered significant at p<10^−50^ and were assigned TF identities by HOMER’s best match to its known vertebrate motif library.

### scATAC-Seq Analysis

UMAP embeddings and TF motif deviation scores for the LaFave et al.^31^ scATAC-Seq dataset were obtained from the interactive Shiny application provided by the original publication, and the underlying data (cell coordinates and motif Z-scores) were downloaded directly from this resource for replotting.

### Tumor Induction Studies

Animal experiments were approved by the MIT Committee on Animal Care. Female NOD *scid* gamma mice (The Jackson Laboratory, 005557) aged 8–12 weeks were used for all experiments. For the tail vein experiment, 1×10^5^ cells were injected in a total volume of 100 μL PBS and mice were euthanized at experiment endpoint (2 weeks following injection). For left ventricle USGI, 5×10^3^ cells were injected in a total volume of 100 μL PBS as previously described^59^. Mice were euthanized at experiment endpoint (∼3 weeks following injection).

The focal lung USGI protocol was developed by Nicole Henning and Laura Maiorino at the Koch Institute. In brief, mice were depilated and imaged by μCT in prone position to obtain three-axis coordinates for injection, including the liver-lung interface, pleural line, and skin. Animals were transferred to the Vevo 3100 ultrasound imager (VisualSonics) and kept in the prone position. The lung parenchyma position was triangulated via the μCT-mediated coordinates, after which 5×10^3^ cells were injected in a total volume of 2 μL PBS, leading to development of a focal nodule in the left lung. Mice were euthanized at experiment endpoint (∼4 weeks following injection).

For the *in vivo* treatment study, 5×10^3^ cells were injected via the tail vein. After allowing the cells to seed for 3 days, tumor seeding was confirmed by intravital bioluminescence imaging (IVIS), and mice were randomized into three treatment groups. Mice were treated daily with either vehicle (0 mg/kg JNJ-64619178), 3 mg/kg JNJ-64619178, or 10 mg/kg JNJ-64619178 in a 20% (2-hydroxypropyl)-β-cyclodextrin (MedChemExpress, HY-101103) solution by oral gavage. Tumor burden was monitored throughout the experiment by intravital bioluminescence imaging (IVIS). Mice were euthanized at experiment endpoint (∼2 weeks).

### Intravital Bioluminescence Imaging (IVIS)

Mice were anesthetized with isoflurane. IVISbrite^TM^ D-Luciferin Potassium Salt (PerkinElmer, 122799) was injected intraperitoneally to reach a concentration of 150 mg/kg body weight. After 8 minutes, mice were placed in the supine position onto the warmed stage in the IVIS Spectrum (PerkinElmer), and isoflurane was continually administered by nose cone. Bioluminescence was detected and measured at 3-minute intervals for 15 minutes using an exposure time of 0.5 seconds to 1 minute, as determined using the auto exposure setting. Total radiance (p/s) was calculated using Living Image v4.7.2.

### Histology and Immunohistochemistry

Tissues were fixed in PBS with 4% formaldehyde and then transferred into 70% ethanol solution. Tissues were processed, embedded in paraffin and sectioned at 4 μm. Following deparaffinization, tissues were stained with Hematoxylin and Eosin, or further processed for IHC. IHC was performed using the LabVision 360 Autostainer with PT Module (Thermo Fisher Scientific). Antigen retrieval was conducted in the PT unit at 97°C for 20 minutes in citrate buffer, pH 6, unless otherwise stated. Sections were stained for 1 hour with antibodies to SDMA (Cell Signaling Technology, 13222S, 1:2,500) and Luciferase (MilliporeSigma, L2164, 1:500, TE Buffer, pH 9).

### Tumor Burden Quantification

Lung tumor burden was assessed by an automated deep neural network developed by Aiforia, in collaboration with the Jacks Lab and Dr. Roderick Bronson. For liver burden, digitized slides were blinded to cell line prior to manual annotation of tumor regions, and tumor areas were computed using QuPath v0.5.0. To calculate percent liver tumor values, liver tumor area was summed and then divided by total liver area. External liver metastases were identified visually and counted. For the *in vivo* treatment study, lung tumor grade was also determined using the Aiforia NSCLC_v2 algorithm. Histological slides were analyzed through consultation with veterinary pathologist Dr. Roderick Bronson.

### ATAC-See Analysis

ATAC-See protocol was adapted from established protocols^64,66,104^. Briefly, fluorescently labeled adapter DNA was prepared by annealing equimolar ME-fw-ATTO550 and ME-rev[Phos] to generate a 45 μM duplex. Tn5 transposome was assembled by incubating equimolar purified Tn5 transposase with adapter duplex for 1 hour at room temperature. Cells were treated with vehicle, 100 nM PRMT5i (JNJ-64619178), 1 μM type I PRMT inhibitor (MS-023), or the combination for 5 days. Cells were harvested by trypsinization. For each reaction, 5×10^5^ cells were washed and permeabilized with NP buffer (0.1% IGEPAL, 0.5% BSA, 1× TAPS), followed by a wash with cold 1× TAPS containing 0.5% BSA. ATAC-See reaction mix was prepared by combining 330 μL nuclease-free water, 100 μL 5× TAPS buffer (50 mM TAPS pH 8.5, 25 mM MgCl_2_), 50 μL DMF, and 20 μL Tn5 transposome. Cells were incubated in 50 μL of the ATAC-See reaction mix at 37°C for 30 minutes. Reactions were stopped with two washes in 100 μL wash buffer (0.1% IGEPAL in 1× PBS). Cells were resuspended in FACS buffer containing DAPI (1 μg/mL final) and incubated on ice for 15 minutes. Flow cytometry was performed on the FACS Celesta HTS-1 (BD Biosciences) and data were analyzed using FlowJo v10.10.0.

### Histone Protein Sequencing

Histone extracts were prepared as described above. Extracts were separated by SDS-PAGE, and gel bands corresponding to histone H2A and histone H4 were excised separately. Proteins were reduced with 20 mM dithiothreitol (MilliporeSigma) for 1 h at 56°C and alkylated with 60 mM iodoacetamide (MilliporeSigma) for 1 h at 25°C in the dark. Proteins were then digested overnight at 37°C with 250 ng α-lytic protease in 100 μL of 100 mM ammonium bicarbonate (pH 8.9).

Peptides were extracted by sequential incubation of gel pieces with 50% acetonitrile/5% formic acid followed by 100 mM ammonium bicarbonate (two rounds), and then 100% acetonitrile followed by 100 mM ammonium bicarbonate (two rounds). All fractions were collected, combined, and reduced to near dryness using a vacuum centrifuge. Samples were desalted using Pierce Peptide Desalting Spin Columns (Thermo Fisher Scientific, 89852) according to the manufacturer’s instructions.

For liquid chromatography-tandem mass spectrometry (LC-MS/MS), peptides were separated by reverse-phase HPLC (Thermo Ultimate 3000) on a PepMap RSLC C18 column (2 μm particle size, 75 μm × 50 cm; Thermo Fisher Scientific, ES903). Solvent A consisted of 0.1% formic acid in water and solvent B of 0.1% formic acid in acetonitrile. Peptides were eluted using the following gradient:

| Time (min) | % B buffer | Flow rate (nL/min) |
| --- | --- | --- |
| 0 | 1 | 300 |
| 20 | 1 | 300 |
| 25 | 1 | 200 |
| 25.5 | 3 | 200 |
| 45 | 23 | 200 |
| 51 | 35 | 200 |
| 53 | 80 | 200 |
| 56 | 80 | 200 |
| 56.1 | 1 | 200 |
| 70 | 1 | 200 |

Eluted peptides were introduced by nano-electrospray ionization into an Orbitrap Exploris 480 mass spectrometer (Thermo Fisher Scientific). The instrument was operated in data-dependent acquisition mode. Full MS scans were acquired at 60,000 resolution over an m/z range of 375– 1500 with a maximum injection time of 25 ms. MS/MS spectra were collected for the top 15 precursor ions using higher-energy collisional dissociation (HCD) at a normalized collision energy of 32 and a resolution of 30,000.

Raw data files (.raw) were analyzed using Sequest HT in Proteome Discoverer (Thermo Fisher Scientific). Search parameters included a precursor mass tolerance of 10 ppm and a fragment mass tolerance of 0.02 Da, allowing up to two missed cleavages for α-lytic protease. Carbamidomethylation of cysteine was set as a fixed modification. Variable modifications included methionine oxidation, methionine loss at the protein N-terminus, N-terminal acetylation, methionine loss plus N-terminal acetylation, arginine methylation, and arginine dimethylation. Data were searched against the UniProt mouse proteome database (UP000000589) along with an in-house contaminant database. Peptide-spectrum matches and peptides were filtered to a 1% false discovery rate (FDR) using Percolator, with a concatenated target-decoy database search strategy and q-value–based validation.

### Immunofluorescence

Cells were seeded onto ethanol-sterilized No. 1.5 glass coverslips and grown to sub-confluence prior to fixation. Coverslips were fixed in 4% paraformaldehyde (PFA) in PBS for 10 min at room temperature, followed by three washes in PBS. Cells were permeabilized with 0.3% Triton X-100 in PBS for 15 min, washed three times with PBS, and blocked for 1 h at room temperature in blocking buffer (5% serum in PBS containing 0.1% Tween-20). Coverslips were then incubated with primary antibodies diluted in blocking buffer for 1 h at room temperature: PRMT5 (Proteintech, 18436-1-AP, 1:100) and PRMT1 (Proteintech, 68365-1-Ig, 1:2,000). Following three PBS washes, samples were incubated with fluorophore-conjugated secondary antibodies diluted in blocking buffer for 1 h at room temperature in the dark: anti-Rabbit IgG (Thermo Scientific, A32733, Alexa Fluor 647, 1:500) and anti-Mouse IgG (Thermo Scientific, A32727, Alexa Fluor 555, 1:500). Slides were also co-stained with CoraLite 488 Phalloidin (Proteintech, PF00001). Coverslips were washed three times with PBS, briefly incubated with DAPI (1:1,000), and mounted onto glass slides using ProLong Gold antifade reagent. Slides were cured overnight at room temperature and sealed prior to imaging. Slides were imaged on a TissueFAXS SL slide scanner (TissueGnostics) equipped with a 40×/0.95 air objective (Zeiss), Spectra III LED light engine (Lumencor), quad-band filter (Chroma Technology set 89402), and sCMOS camera (Hamamatsu ORCA-Fusion BT).

For quantification, image analysis was performed in R v4.3.2 using the EBImage package (v4.44.0)^105^. Nuclear segmentation was performed on the DAPI channel. Images were smoothed with a Gaussian filter (σ=2 pixels, 0.325 μm) and binarized using Otsu’s method. Binary masks were refined by hole-filling and morphological opening with a 5-pixel disc structuring element. Touching nuclei were separated by a distance-transform watershed: a Euclidean distance map was computed from the binary mask, local maxima were identified within a 15-pixel radius, and seeds were expanded back into the binary mask. Objects smaller than 200 pixels or larger than 20,000 pixels were excluded as debris or segmentation artifacts. Cytoplasmic signal was sampled from a fixed-width annular ring of 15 pixels (2.44 μm) surrounding each nuclear mask, clipped to that cell’s Voronoi territory to prevent overlap with neighboring nuclei. Per-channel background was estimated as the second percentile of pixel intensities in regions outside all cell masks and subtracted from nuclear and cytoplasmic means prior to ratio calculation. Cells in which the background-corrected cytoplasmic mean was less than twice the background standard deviation were excluded from ratio calculations for that channel. The nuclear-to-cytoplasmic ratio was expressed as log_2_(nuclear mean/cytoplasmic mean), such that values greater than zero indicate nuclear enrichment, values less than zero indicate cytoplasmic enrichment, and zero indicates equal distribution between compartments. Portions of the image analysis code were developed with the assistance of Claude Sonnet 5 (Anthropic); all code was reviewed, validated, and executed by the authors.

### TCGA Patient Analysis

Processed RNA-Seq data for TCGA Firehose Legacy and TCGA PanCancer^106^ was obtained from cBioPortal^107–109^. Patient samples were segregated on *CDKN2A* status in cBioPortal with the following command:

~~~
CDKN2A: HOMDEL
~~~

### Statistical Analysis and Plotting Software

R v4.3.2 and GraphPad Prism v10.1.0 were used to perform statistical analyses. R v4.3.2, GraphPad Prism v10.1.0, and Adobe Illustrator v28.2 were used for generating and plotting figures. All statistical tests are two-sided where applicable. Differences were considered statistically significant if p<0.05, while those with p≥0.05 were considered not significant (n.s.).

## Acknowledgments

We thank the Koch Institute’s Robert A. Swanson (1969) Biotechnology Center for technical support, particularly the Hope Babette Tang (1983) Histology, Barbara K. Ostrom (1978) Integrated Genomics and Bioinformatics, Preclinical Modeling, Imaging & Testing, Biopolymers & Proteomics, and Microscopy Core Facilities. Images in schematics were created using BioRender.

## Author Contributions

CEF and JAL conceived this study with input from HSM, AAB, TJ, YSF, and LML. CEF conducted all of the experiments with assistance from: NAO and RH for *in vivo* studies and cell line characterization; AG for chromVAR analysis; ZY for ATAC-See experimentation and analysis; NMA for Aiforia analysis; and GJS for preparation of ATAC-Seq samples. NH performed all injections. CEF, NAO, and JAL interpreted the data. CEF and JAL wrote the manuscript, with input from other authors.

## Data and Material Availability

All data needed to evaluate the conclusions in the paper are present in the paper and/or the Supplemental Figures. RNA-Seq and ATAC-Seq data are being deposited in the Gene Expression Omnibus (GEO). Mass spectrometry data are being deposited to the ProteomeXchange Consortium via the PRIDE partner repository. Accession codes and reviewer tokens will be made available when submissions are approved. All data will be made public upon publication. Correspondence and all requests should be sent to Jacqueline A. Lees.

