## Supplemental Figures and Legends for "PRMT5 inhibition induces a histone methylation switch that drives chromatin plasticity, drug resistance, and metastatic progression"

[1] The David H. Koch Institute for Integrative Cancer Research, Massachusetts Institute of Technology, Cambridge, MA, USA
[2] Department of Biology, Massachusetts Institute of Technology, Cambridge, MA, USA
[3] Department of Cell Biology, Albert Einstein College of Medicine, Bronx, NY, USA
[4] Montefiore Einstein Comprehensive Cancer Center, Albert Einstein College of Medicine, Bronx, NY, USA
[5] Molecular Pharmacology Program, Sloan Kettering Institute, Memorial Sloan Kettering Cancer Center, New York, NY, USA
[6] Ruth L. and David S. Gottesman Institute for Stem Cell Biology and Regenerative Medicine, Albert Einstein College of Medicine, Bronx, NY, USA
[7] Cancer Dormancy & Tumor Microenvironment Institute, Albert Einstein College of Medicine, Bronx, NY, USA
[*] Corresponding Author:
Jacqueline A. Lees
The David H. Koch Institute for Integrative Cancer Research & Department of Biology
Massachusetts Institute of Technology
500 Main Street, 76-261C, Cambridge, MA, 02139


### Supplemental Figure 1

**A**

| Cell Line | Prior Nomenclature |
|---|---|
| KP^Early-Parental | KP2 (Ref. 28), KP1233 (Ref. 95) |
| KP^Early-Resistant (R1, R2, R3) | KP2-R7, R1, R4 (Ref. 28) |
| KP^Late-Parental | KP1 (Ref. 28), KP393T5 (Ref. 29) |
| KP^Late-Resistant (R1, R2, R3) | KP1-R6, R3, R7 (Ref. 28) |

Resistant variant used is Resistant 1, unless otherwise noted

**B**

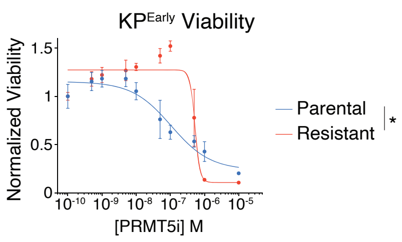


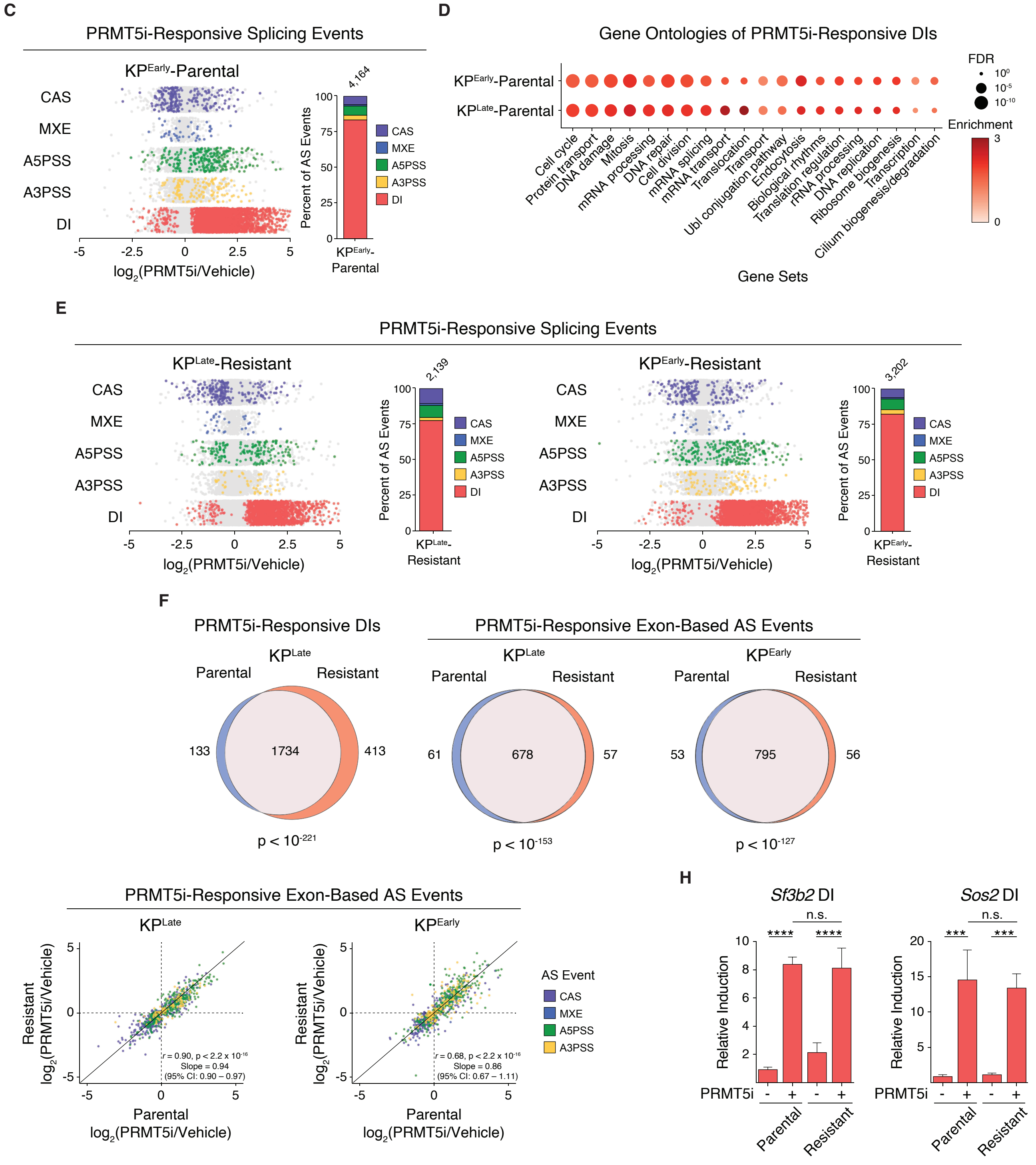

**Supplemental Figure 1. PRMT5i-induced AS is equal in parental and resistant lines.**
**A.** Cell lines used in this study, including prior names and publications.
**B.** Relative viability of $KP^{Early}$-Parental (blue) and $KP^{Early}$-Resistant (red) after 5-day treatment with the indicated JNJ-64619178 (PRMT5i) concentrations. Data are mean ± SD of 3 technical replicates per line. Significance compares area under the curve (AUC) by Welch's t-test.
**C.** $KP^{Early}$-Parental cells were treated with vehicle or 100 nM JNJ-64619178 (PRMT5i) for 3 days and poly(A)+ mRNA deep sequenced to identify and quantify DIs, as well as exon-based alternative splicing (AS) events: cassette exons (CAS), mutually exclusive exons (MXE), and alternative 5′ or 3′ splice sites (A5PSS, A3PSS). (*Left*) Dots represent individual splicing events and denote those significantly ($p_{adj}<0.05$, color), or not significantly (gray), altered by treatment. (*Right*) The graph indicates the proportion of each splicing type that is significantly ($p_{adj}<0.05$) altered by PRMT5i. The total number of significant splicing events is indicated above.
**D.** Enriched GO terms from the KW Biological Process Gene Set for DIs that are significantly altered ($p_{adj}<0.05$) by 3-day PRMT5i treatment of $KP^{Early}$- and $KP^{Late}$-Parental cells. Terms are displayed if they are significant in at least one cell line (FDR<0.01). Color represents enrichment score and point size is scaled inversely to FDR, such that larger points indicate greater significance.
**E.** Cells were treated for 3 days with vehicle or PRMT5i (10 µM EPZ015666 for $KP^{Late}$-Resistant and 100 nM JNJ-64619178 for $KP^{Early}$-Resistant) and poly(A)+ mRNA deep sequenced to identify and quantify DIs and exon-based AS events. For both resistant lines: (*left*) the dots represent individual events for the indicated splicing types and denote those significantly ($p_{adj}<0.05$, color), or not significantly (gray), altered by PRMT5i; (*right*) the graph shows the proportion of each splicing type that is significantly ($p_{adj}<0.05$) altered by PRMT5i, with the total number of significant events indicated above.
**F.** Overlap of DIs, or exon-based AS events, that are significantly induced (PRMT5i-responsive) by 3-day PRMT5i treatment of parental and resistant $KP^{Late}$ (10 µM EPZ015666) and $KP^{Early}$ (100 nM JNJ-64619178). Significance determined by hypergeometric test.
**G.** Relative induction of exon-based AS events in parental and resistant lines for $KP^{Early}$ (*left*) and $KP^{Late}$ (*right*) after a 3-day PRMT5i treatment. Solid line indicates a line of identity. Pearson correlation and Deming regression values are displayed, indicating similar AS induction regardless of resistance status.
**H.** qRT-PCR quantification of DI levels for the indicated genes (*Sf3b2* and *Sos2*) in $KP^{Late}$-Parental and Resistant lines after a 3-day PRMT5i treatment (10 µM EPZ015666). DI levels were normalized to total gene levels. Two-way ANOVA, with treatment and resistance status as main effects, was performed on the normalized DI levels. Data are mean ± SD of 3 technical replicates per condition.
For all panels, * indicates p<0.05, *** indicates p<0.001, **** indicates p<0.0001, and n.s. indicates not significant.

### Supplemental Figure 2

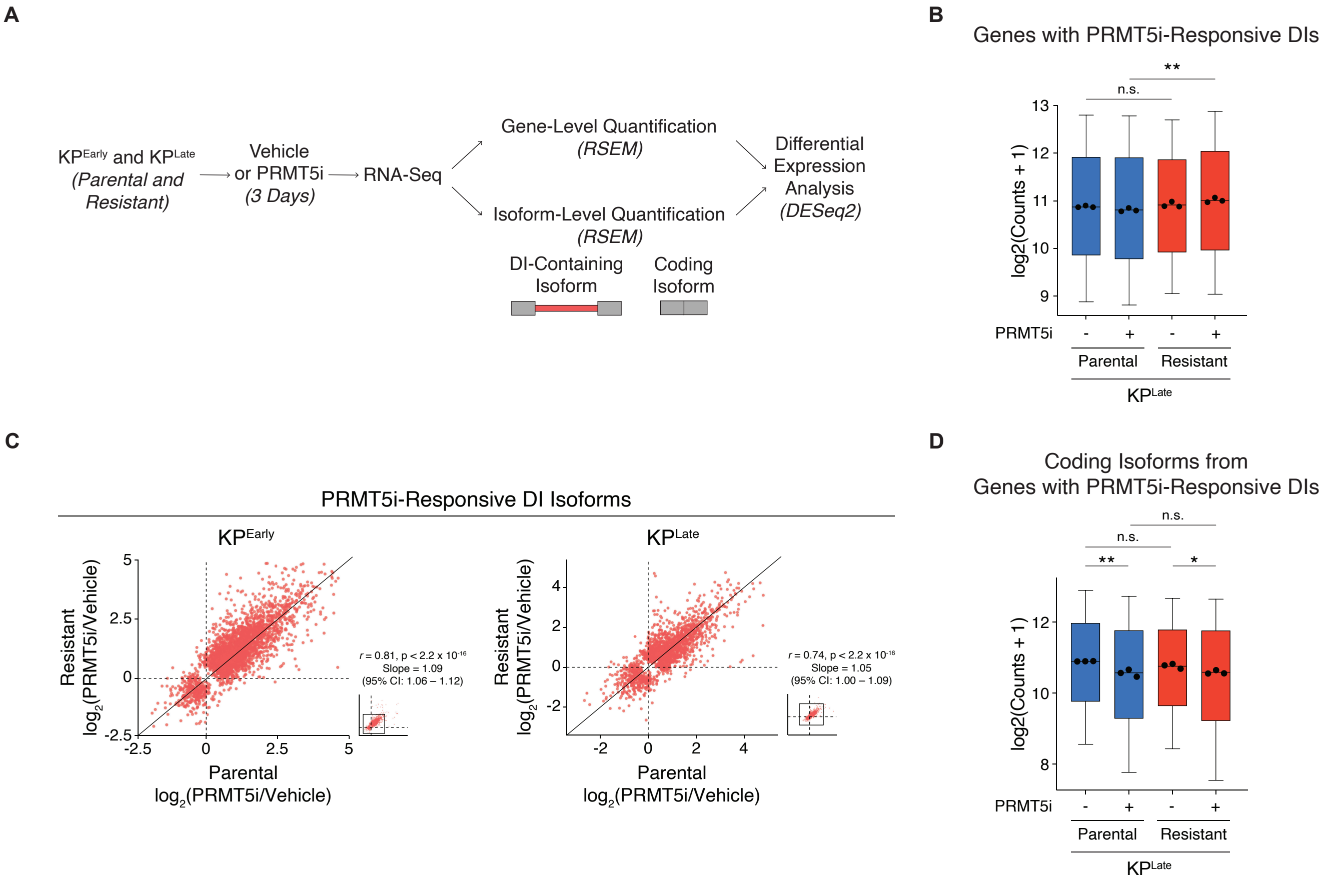

**Supplemental Figure 2. Coding isoforms of DI-containing genes are equally downregulated in parental and resistant lines upon PRMT5 inhibition.**

**A.** Experimental schematic for gene- and isoform-level (including coding and DI isoforms) quantification and differential expression analysis.

**B.** Normalized expression counts of genes with PRMT5i-responsive DIs in $KP^{Late}$-Parental (blue) and Resistant (red) lines after 3-day vehicle or PRMT5i treatment. Box plots were generated from average $log_2$(Counts+1) for each gene (n=1,560), with individual points overlaid representing median counts for each replicate (n=3).

**C.** Relative induction of DI isoforms in parental and resistant lines for $KP^{Early}$ (*left*) and $KP^{Late}$ (*right*) after 3-day PRMT5i treatment. Solid line indicates a line of identity. Inset shows the full data range, with the boxed region indicating the axis limits of the main plot. Pearson correlation and Deming regression values are displayed.

**D.** Normalized expression counts of coding isoforms from genes with PRMT5i-responsive DIs in $KP^{Late}$-Parental (blue) and Resistant (red) lines after 3-day vehicle or PRMT5i treatment. Box plots were generated from average $log_2$(Counts+1) for each coding isoform (n=1,560), with individual points overlaid representing median counts for each replicate (n=3).

For corresponding panels, two-way ANOVA, with treatment and resistance status as main effects, was performed on the median counts for each replicate. * indicates $p<0.05$, ** indicates $p<0.01$, and n.s. indicates not significant.

### Supplemental Figure 3

**A**

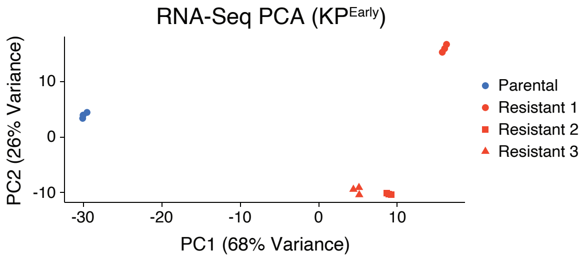


**B**

| Abbreviation | MSigDB Gene Set |
|---|---|
| EMT Up | GO MESENCHYME DEVELOPMENT |
| EMT Down | GOTZMANN EPITHELIAL TO MESENCHYMAL TRANSITION DN |
| Progression Up | VECCHI GASTRIC CANCER ADVANCED VS EARLY UP |
| Progression Down | VECCHI GASTRIC CANCER ADVANCED VS EARLY DOWN |
| Metastasis Up | TOMLINS METASTASIS UP |
| Metastasis Down | JAEGER METASTASIS DOWN |

**C**

Normalized Expression Counts (RNA-Seq)

| | | $KP^{Early}$ | | | | $KP^{Late}$ | | | |
|---|---|---|---|---|---|---|---|---|---|
| | | | Resistant | | | | Resistant | | |
| | | Parental | 1 | 2 | 3 | Parental | 1 | 2 | 3 |
| Squamous | *Krt5* | 0 | 0 | 0 | 0 | 0 | 0 | 1 | 0 |
| | *Krt6a* | 0 | 0 | 0 | 0 | 37 | 4 | 2 | 1 |
| | *Krt14* | 74 | 29 | 24 | 19 | 9 | 7 | 1 | 1 |
| Neuroendocrine | *Ascl1* | 0 | 0 | 0 | 0 | 1 | 0 | 0 | 0 |
| | *Chga* | 0 | 0 | 0 | 0 | 0 | 0 | 0 | 0 |
| | *Scg3* | 0 | 0 | 0 | 0 | 0 | 0 | 0 | 0 |

**D**

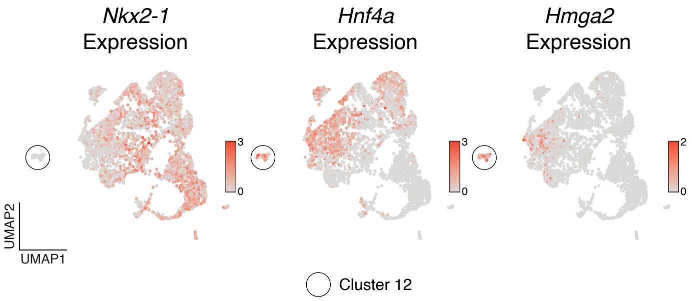


**E**

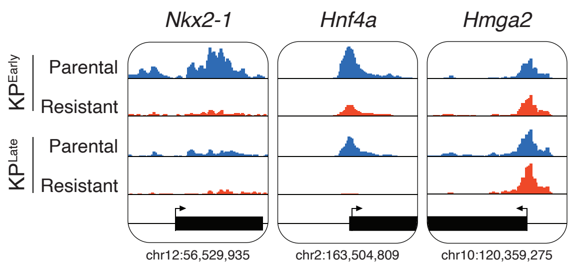


**F**

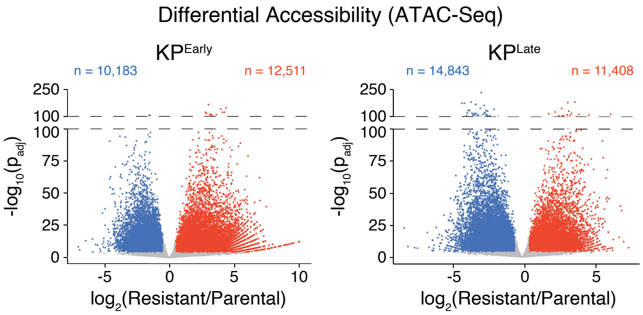


**G**

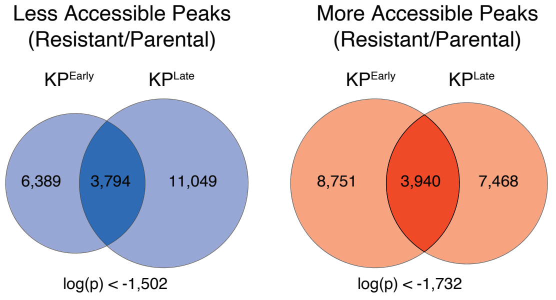


**H**

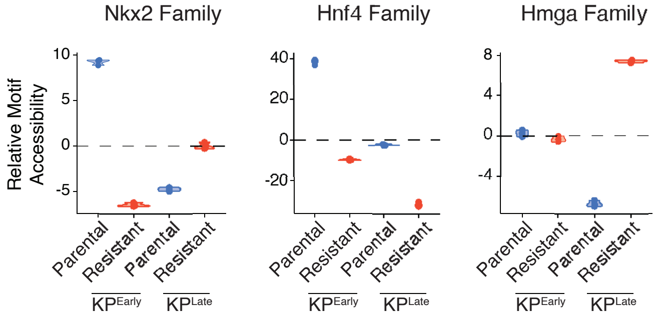

**Supplemental Figure 3. PRMT5 inhibitor resistance induces broad chromatin accessibility changes that reflect naturally progressed disease.**

**A.** PCA of the 500 most variably expressed genes by RNA-Seq in $KP^{Early}$-Parental (blue) versus three independently derived $KP^{Early}$-Resistant populations (red) with triplicate samples for each.

**B.** Table of abbreviated and full MSigDB gene set names for gene sets plotted in **Fig. 2C**.

**C.** Table of normalized RNA counts for the indicated squamous or neuroendocrine transdifferentiation markers in $KP^{Early}$- and $KP^{Late}$-Parental and Resistant lines. Data are mean of 3 biological replicates per line.

**D.** Expression of dedifferentiation markers in scRNA-Seq UMAP from an autochthonous model of KP-mutant LUAD.

**E.** Genomic accessibility of the indicated dedifferentiation and resistance markers in $KP^{Early}$- and $KP^{Late}$-Parental (blue) and Resistant (red) lines.

**F.** Differential chromatin accessibility comparing (*left*) $KP^{Early}$- and (*right*) $KP^{Late}$-Parental to their respective Resistant lines. Points represent individual loci, where red indicates more accessible ($p_{adj}<10^{-4}$, $log_2FC>0.5$), blue indicates less accessible ($p_{adj}<10^{-4}$, $log_2FC<-0.5$), and gray indicates not differentially accessible.

**G.** Venn diagrams show the number of peaks that are significantly (*left*) less accessible or (*right*) more accessible in resistant versus parental lines for $KP^{Early}$ and $KP^{Late}$, with the significance of the overlap between the two cell lines shown below the intersection.

**H.** chromVAR deviation scores for combined (*left*) Nkx2, (*center*) Hnf4, and (*right*) Hmga family motifs in $KP^{Early}$- and $KP^{Late}$-Parental (blue) and Resistant (red) lines. Each point represents one biological replicate (n=3).

### Supplemental Figure 4

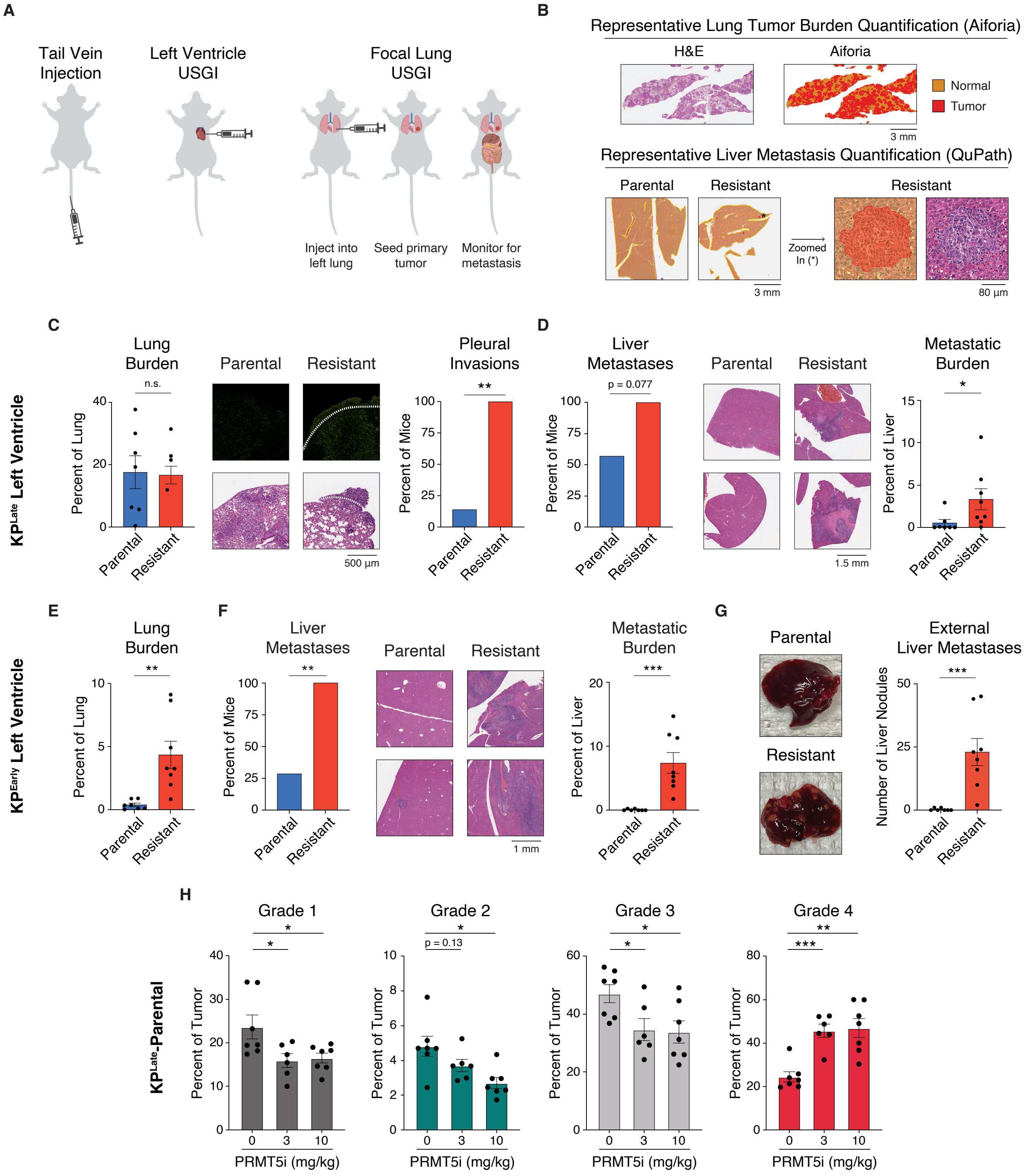

**Supplemental Figure 4. PRMT5i-resistant cells are hyper-metastatic in multiple *in vivo* models, and PRMT5i treatment of parental tumors increases tumor grade.**

**A.** Schematics of (*left*) tail vein injection, (*center*) left ventricle ultrasound-guided injection (USGI), and (*right*) focal lung USGI.

**B.** (*Top*) Representative Aiforia-generated images of lung tumor burden quantification for **Fig. 3A**. (*Bottom*) Representative QuPath-generated images of liver metastasis burden quantification for **Fig. 3B**. For both, normal tissue is shown in orange and tumor is shown in red.

**C–D.** Left ventricle USGI was used to introduce $5\times10^3$ $KP^{Late}$-Parental or Resistant cells into 7–8 recipients per line. These were analyzed for: **(C)** lung tumors, with tumor burden (p=0.96), representative H&E images, and pleural invasion quantification (p=0.0014) shown; and **(D)** liver metastases, with metastatic frequency (p=0.077), representative H&E images, and burden (p=0.014) shown.

**E–G.** Left ventricle USGI was used to introduce $5\times10^3$ $KP^{Early}$-Parental or Resistant cells into 7–8 recipients per line. These were analyzed for: **(E)** lung tumor burden (p=0.0012), **(F)** liver metastases, with metastatic frequency (p=0.0070), representative H&E images, and burden (p=0.0003) shown, and (**G**) external liver metastases, with representative images of livers and quantification of external nodules (p=0.0003) shown.

**H.** Component parts of tumor grade analysis for **Fig. 3I** (n=6–7 recipients per condition).

For all panels, data are mean ± SEM where applicable. Fisher's exact test was used for frequency of tumors, metastases, and pleural invasions, Mann-Whitney U-test was used for burden, and Welch's t-test was used for *in vivo* treatment grade. * indicates p<0.05, ** indicates p<0.01, *** indicates p<0.001, and n.s. indicates not significant.

**Supplemental Figure 5**

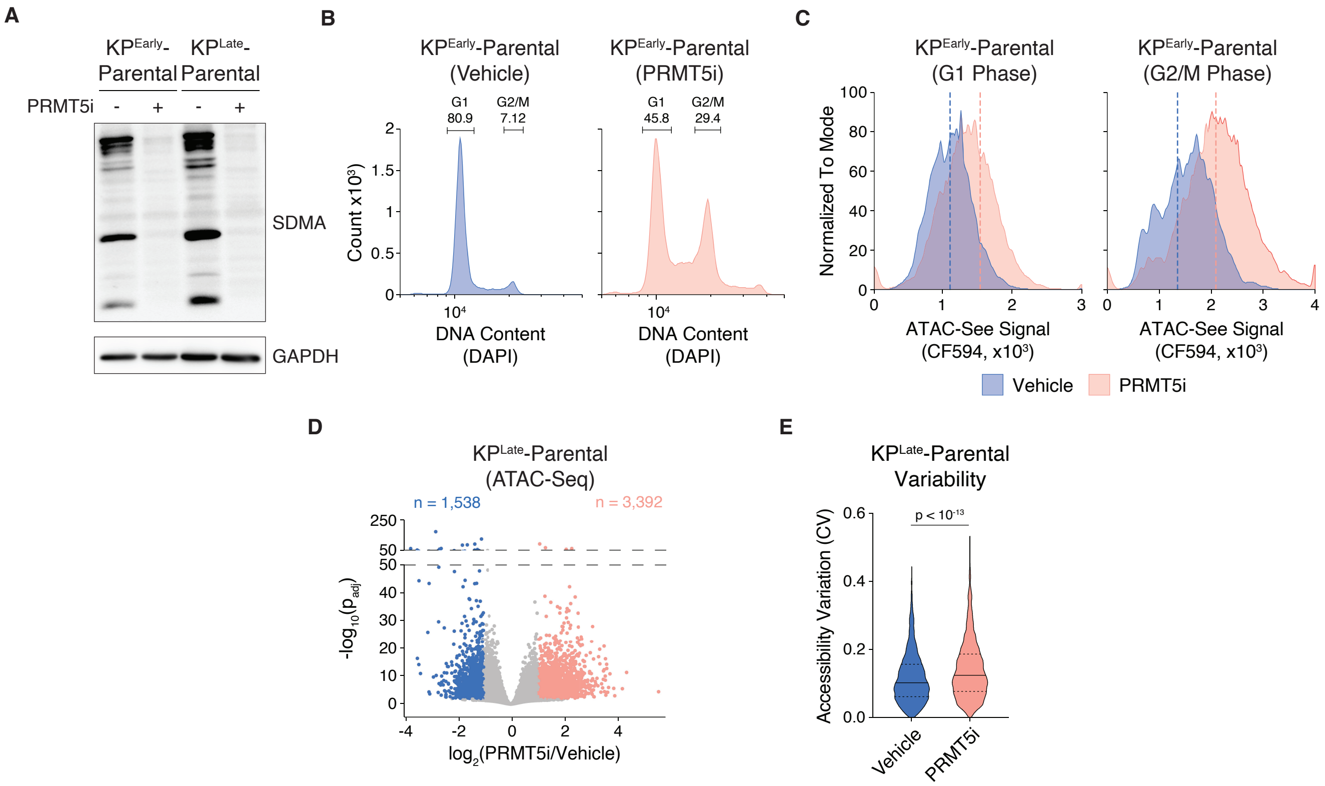

**Supplemental Figure 5. PRMT5 inhibition induces broad chromatin accessibility changes.**

**A.** Western blot analysis of symmetric dimethylarginine (SDMA) levels in $KP^{Early}$- and $KP^{Late}$-Parental cells treated with vehicle or PRMT5i (100 nM JNJ-64619178) for 5 days. GAPDH is shown as a loading control.

**B.** Flow cytometry analysis of DNA content by DAPI staining from representative ATAC-See samples after 5-day vehicle or PRMT5i treatment. ATAC-See signal was quantified from the indicated gates corresponding to G1 and G2/M populations.

**C.** Representative ATAC-See signal distributions for $KP^{Early}$-Parental cells following 5-day vehicle or PRMT5i treatment. Dashed lines indicate the mean fluorescence intensity (MFI) for each sample.

**D.** Differential chromatin accessibility in $KP^{Late}$-Parental cells after 5-day vehicle or PRMT5i treatment. Each point represents an individual genomic locus that is more accessible (pink, $p_{adj}<0.01$, $\log_2FC>1$), less accessible (blue, $p_{adj}<0.01$, $\log_2FC<-1$), or not significantly changed (gray).

**E.** Coefficient of variation (CV) across replicates for loci with increased accessibility in $KP^{Late}$-Parental cells after 5-day vehicle (blue) or PRMT5i (pink) treatment. Wilcoxon rank-sum test.

**Supplemental Figure 6**

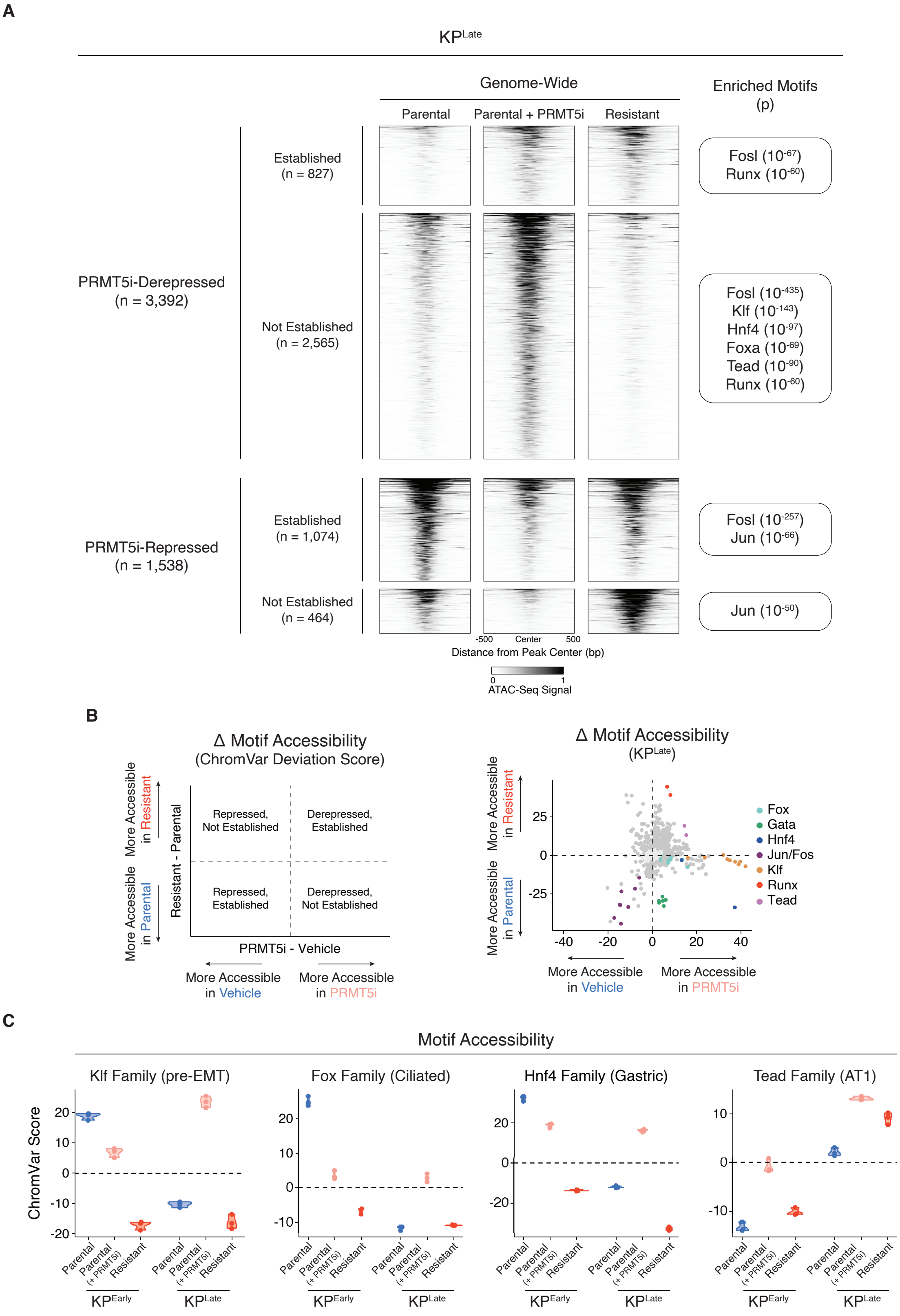

**Supplemental Figure 6. Chromatin accessibility programs induced by PRMT5 inhibition largely map to canonical KP-mutant LUAD dedifferentiation states.**

**A.** Genome-wide analysis of chromatin accessibility changes following 5-day PRMT5 inhibition in $KP^{Late}$-Parental cells and their relationship to accessibility states established in resistant cells. Four categories of loci are shown based on whether PRMT5i induces derepression or repression of accessibility and whether these changes are established or not established in the resistant state: derepressed-and-established, derepressed-and-not established, repressed-and-established, and repressed-and-not established. (*Left*) Genome-wide accessibility heatmaps. (*Right*) Enriched *de novo* TF motifs for each category, labeled by best-matching TF ($p<10^{-50}$).

**B.** chromVAR was conducted for 5-day vehicle- and PRMT5i-treated $KP^{Late}$-Parental cells (x-axis) and resistant versus parental $KP^{Late}$ cells (y-axis). (*Left*) Schematic illustrating whether motif accessibility changes represent short-term derepression or repression and whether these changes are established or not established in the resistant state. (*Right*) Deviation scores, where individual points represent motifs that are significantly (dark gray or colored, $p_{adj}<0.01$) or non-significantly (light gray) altered. Colored points indicate motifs belonging to the same TF family.

**C.** Individual chromVAR deviation scores for selected TF family motifs in $KP^{Early}$ and $KP^{Late}$ showing accessibility in 5-day vehicle (blue) or PRMT5i (pink) treated parental cells and stably resistant cells (red). Each point represents one biological replicate (n=3).

**Supplemental Figure 7**

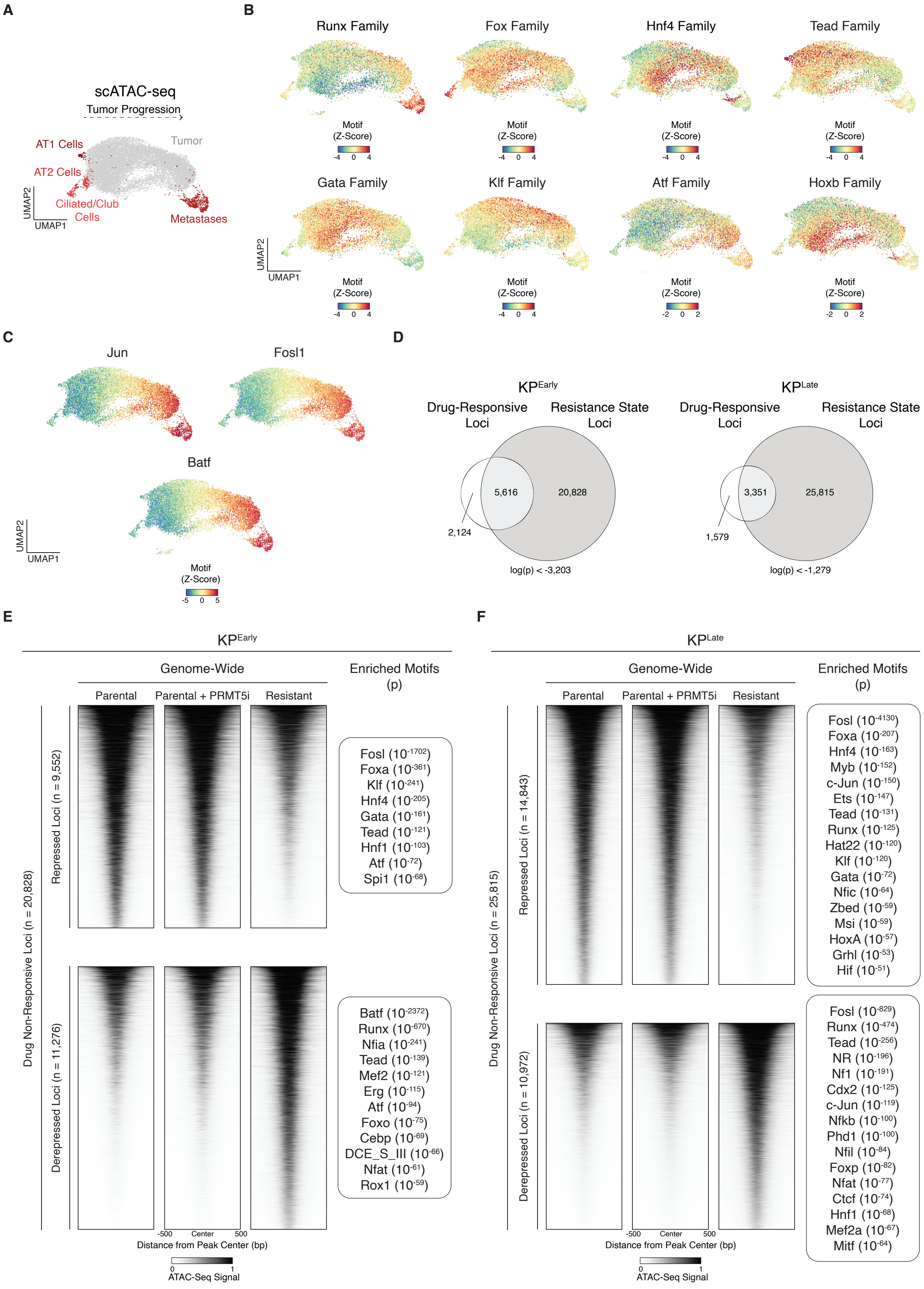

**Supplemental Figure 7. Chromatin accessibility programs induced by PRMT5 inhibition are selectively established in resistant cells.**

**A–C.** Chromatin accessibility and TF motif activity along tumor progression in an autochthonous mouse model of KP-mutant LUAD. (**A**) UMAP projection of scATAC-Seq profiles from normal and neoplastic lung, ordered along a continuum of tumor progression[31]. Points represent single cells colored by major annotated populations, including AT1 cells, AT2 cells, ciliated/club cells, primary tumor cells (gray), and metastatic cancer cells. (**B–C**) UMAPs of motif accessibility Z-scores for the indicated TF motifs: (**B**) shows TF motifs that exhibit enrichment in specific KP-mutant LUAD states; and (**C**) shows AP-1 TFs, including Jun, Fosl1, and Batf. For all motif plots, colors indicate relative motif accessibility (Z-score; blue, low; red, high).

**D.** Overlap between loci that become differentially accessible after 5-day PRMT5i treatment (drug-responsive peaks) and those that are differentially accessible in resistant cells (resistance-state peaks) for (*left*) $KP^{Early}$ and (*right*) $KP^{Late}$. Hypergeometric test.

**E–F.** Genome-wide analysis of chromatin accessibility loci that are unchanged following 5-day PRMT5 inhibition but are differentially accessible in resistant cells for: (**E**) $KP^{Early}$ and (**F**) $KP^{Late}$ cells. Two categories of loci are shown based on whether accessibility is lost or gained in the resistant state. (*Left*) Genome-wide accessibility heatmaps. (*Right*) Enriched *de novo* TF motifs for each category, labeled by best-matching TF ($p<10^{-50}$).

### Supplemental Figure 8

**A**

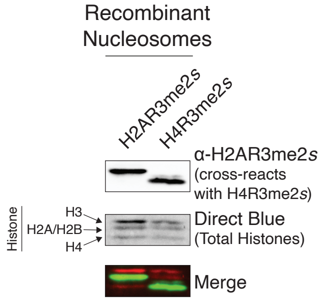


**B**

**Histone H2A Peptides**

| Sequence | Residues | Modifications | Total Spectra |
|---|---|---|---|
| MSGRGKQGGKA | 1-11 | Met-loss+Acetyl; R4(Dimethyl) | 6 |
| RDNKKTRIIPRHLQLA | 72-87 | | 2 |
| RIIPRHLQLA | 77-87 | | 328 |
| IRNDEELNKLLGKV | 88-101 | | 17 |
| IRNDEELNKLLGKVT | 88-102 | | 30 |
| QGGVLPNIQA | 105-114 | | 8 |
| QGGVLPNIQAV | 105-115 | | 28 |

**C**

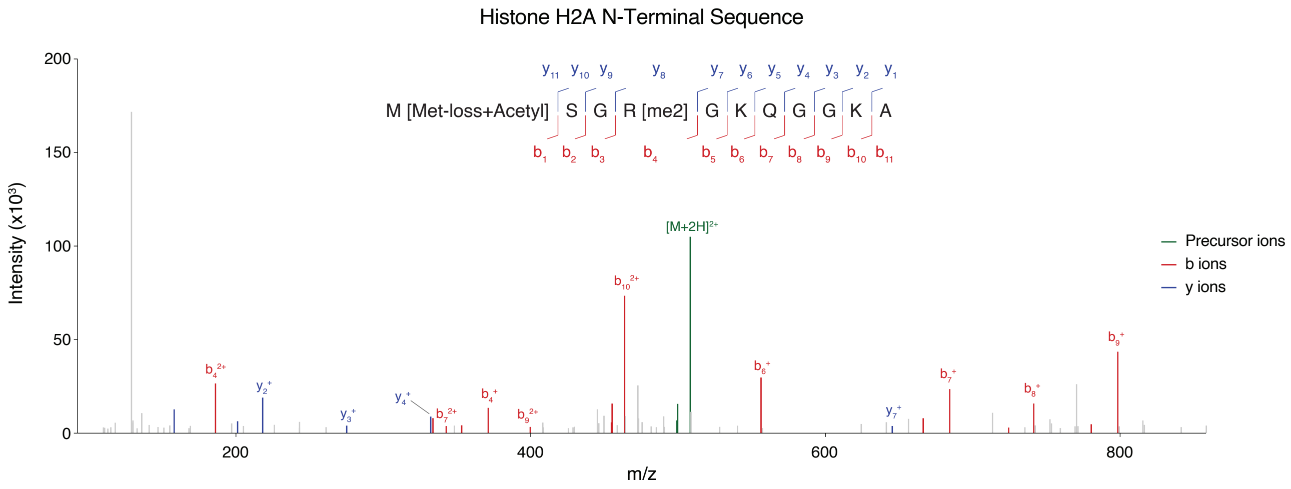


**D**

**Histone H4 Peptides**

| Sequence | Residues | Modifications | Total Spectra |
|---|---|---|---|
| MSGRGKGGKGLGKGGA | 1-16 | Met-loss+Acetyl | 6 |
| LRDNIQGIT | 23-31 | | 20 |
| LRDNIQGITKPA | 23-34 | | 61 |
| YALKRQGRTLYGFGG | 89-103 | | 25 |
| LKRQGRTLYGFGG | 91-103 | | 5 |

**E**

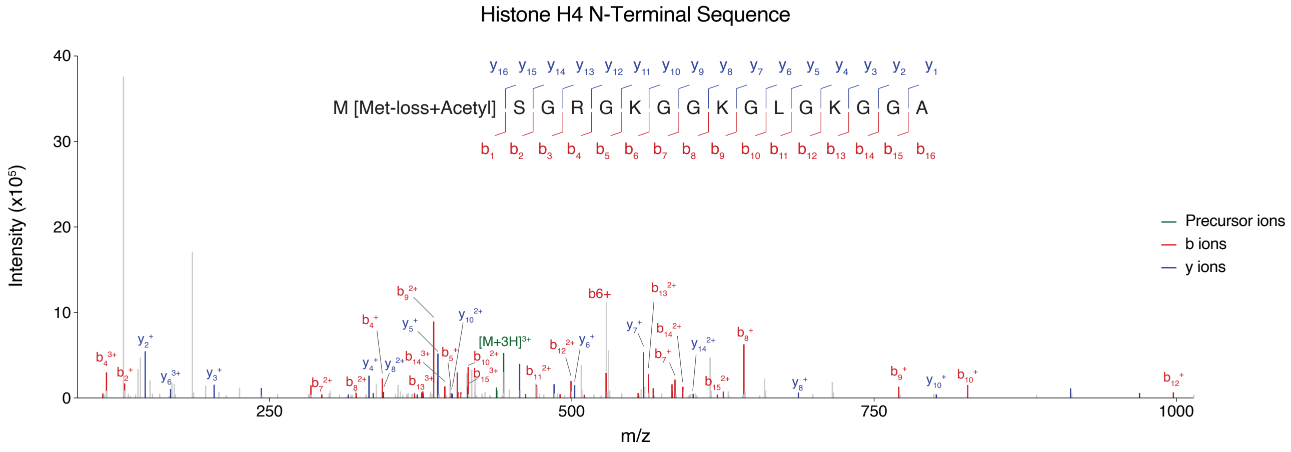

**Supplemental Figure 8. Arginine dimethylation is detected on histone H2A but not H4.**
**A.** Western blot of recombinant nucleosomes containing either H2AR3me2*s* or H4R3me2*s*, probed with an α-H2AR3me2*s* antibody. A band is detected in both lanes, indicating that the antibody cross-reacts with the R3me2*s* mark on H4. Direct Blue staining confirms total histone loading, and the merged fluorescence image confirms the identity of each modified histone based on molecular weight.
**B.** Table of peptides identified by LC-MS/MS mapping to histone H2A from $KP^{Late}$-Parental cells, including primary sequence, residue numbers, post-translational modifications, and spectral counts.
**C.** MS2 spectrum of the N-terminal H2A peptide. The identified sequence and modifications (bracketed) are shown at the top with annotated b and y fragment ions. Precursor ions are shown in green, while b and y ions are shown in red and blue, respectively.
**D.** Table of peptides identified by LC-MS/MS mapping to histone H4 from $KP^{Late}$-Parental cells, including primary sequence, residue numbers, post-translational modifications, and spectral counts.
**E.** MS2 spectrum of the N-terminal H4 peptide. The identified sequence and modifications (bracketed) are shown at the top with annotated b and y fragment ions. Precursor ions are shown in green, while b and y ions are shown in red and blue, respectively.

**Supplemental Figure 9**

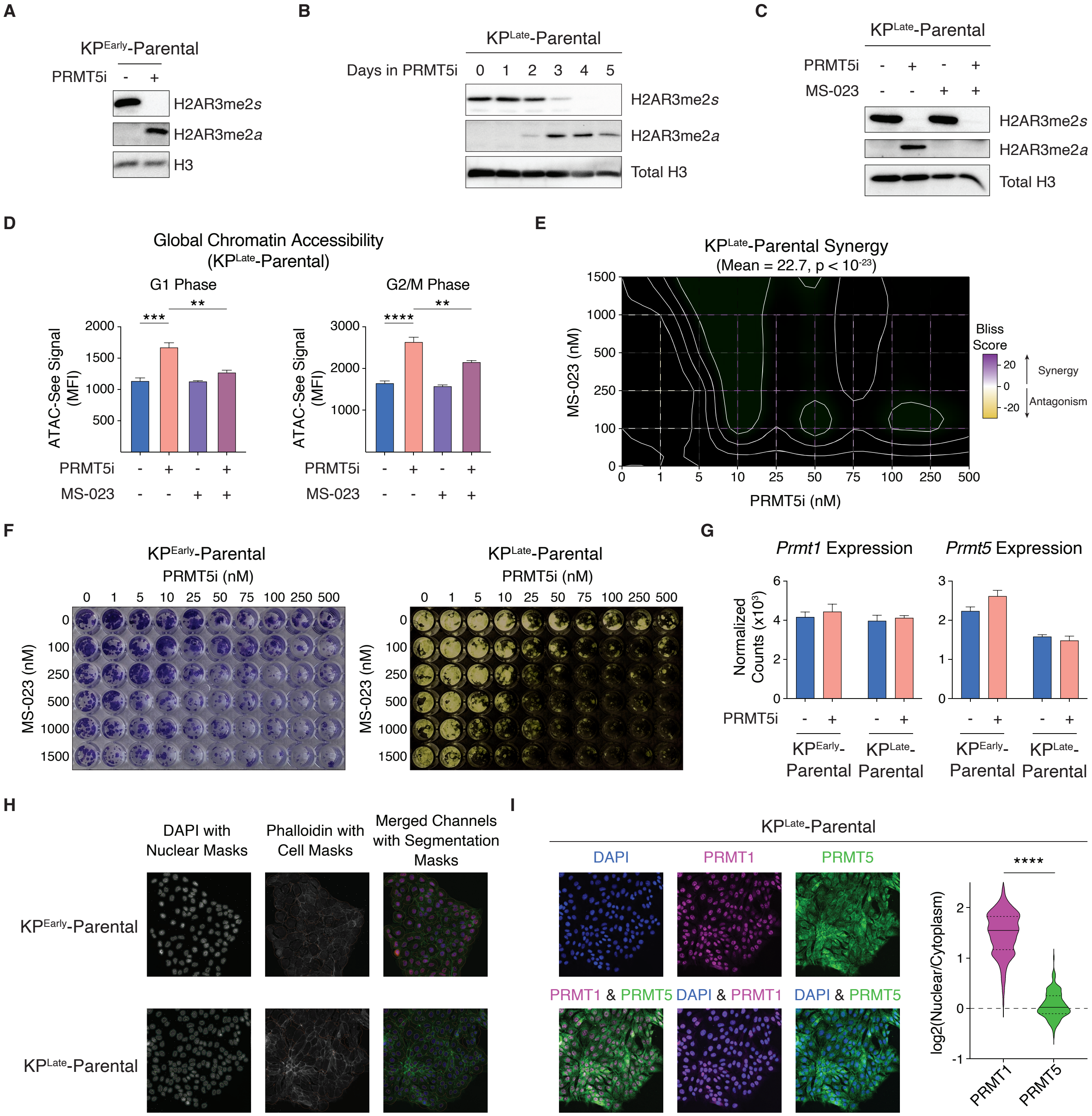

**Supplemental Figure 9. PRMT5 gatekeeps a symmetric-to-asymmetric H2AR3 methylation switch.**

**A.** Western blot of H2AR3me2*s* and H2AR3me2*a* in $KP^{Early}$-Parental cells after a 5-day vehicle or PRMT5i (100 nM JNJ-64619178) treatment. Total H3 serves as a loading control.

**B.** Western blot of H2AR3me2*s* and H2AR3me2*a* in $KP^{Late}$-Parental cells during a 5-day time course of PRMT5i treatment. Total H3 serves as a loading control.

**C.** Western blot of H2AR3me2*s* and H2AR3me2*a* in $KP^{Late}$-Parental cells after a 5-day vehicle, PRMT5i, MS-023, or combination treatment. Total H3 serves as a loading control.

**D.** Mean fluorescence intensity (MFI) of ATAC-See signal in $KP^{Late}$-Parental cells treated with vehicle, 100 nM PRMT5i, MS-023, or the combination for 5 days. G1 and G2/M populations were considered separately. Data are mean ± SD of 3 biological replicates per condition.

**E.** Bliss synergy contour map for 5-day co-treatment of $KP^{Late}$-Parental cells with various PRMT5i (x-axis) and MS-023 (y-axis) concentrations. Color indicates the Bliss synergy score, with yellow indicating antagonism and purple indicating synergy.

**F.** Representative images of crystal violet staining used for synergy analysis in (*left*) $KP^{Early}$- and (*right*) $KP^{Late}$-Parental. PRMT5i doses are indicated for each column and MS-023 doses are indicated for each row.

**G.** Relative expression of (*left*) *Prmt1* and (*right*) *Prmt5* mRNA in $KP^{Early}$- and $KP^{Late}$-Parental cells upon 3-day vehicle (blue) or PRMT5i (pink) treatment. Data are mean ± SD of 3 biological replicates per line.

**H.** Representative immunofluorescence images of $KP^{Early}$- and $KP^{Late}$-Parental cells used for PRMT1 and PRMT5 subcellular localization. (*Left*) DAPI staining was used to generate nuclear masks, which served as seeds for Voronoi cell territories. (*Middle*) Phalloidin staining is shown to illustrate cell boundaries; Voronoi territories were derived from the DAPI-seeded nuclear masks rather than from the phalloidin signal. (*Right*) For nuclear-to-cytoplasmic ratio measurements (**Fig. 5H** and **Fig. S9I**), the cytoplasmic region was defined as a 15-pixel-wide annulus surrounding each DAPI-labeled nucleus and clipped to the corresponding Voronoi territory.

**I.** Localization of PRMT1 and PRMT5 in $KP^{Late}$-Parental cells. (*Left*) Immunofluorescence images of endogenous PRMT1 (magenta), PRMT5 (green), and DAPI (blue), with merged views highlighting nuclear enrichment of PRMT1. (*Right*) Violin plots depict single-cell $\log_2$(nuclear/cytoplasmic) intensity ratios (n=162 cells).

For corresponding panels, Welch's t-test was used. ** indicates $p<0.01$, *** indicates $p<0.001$, and **** indicates $p<0.0001$.

**Supplemental Figure 10**

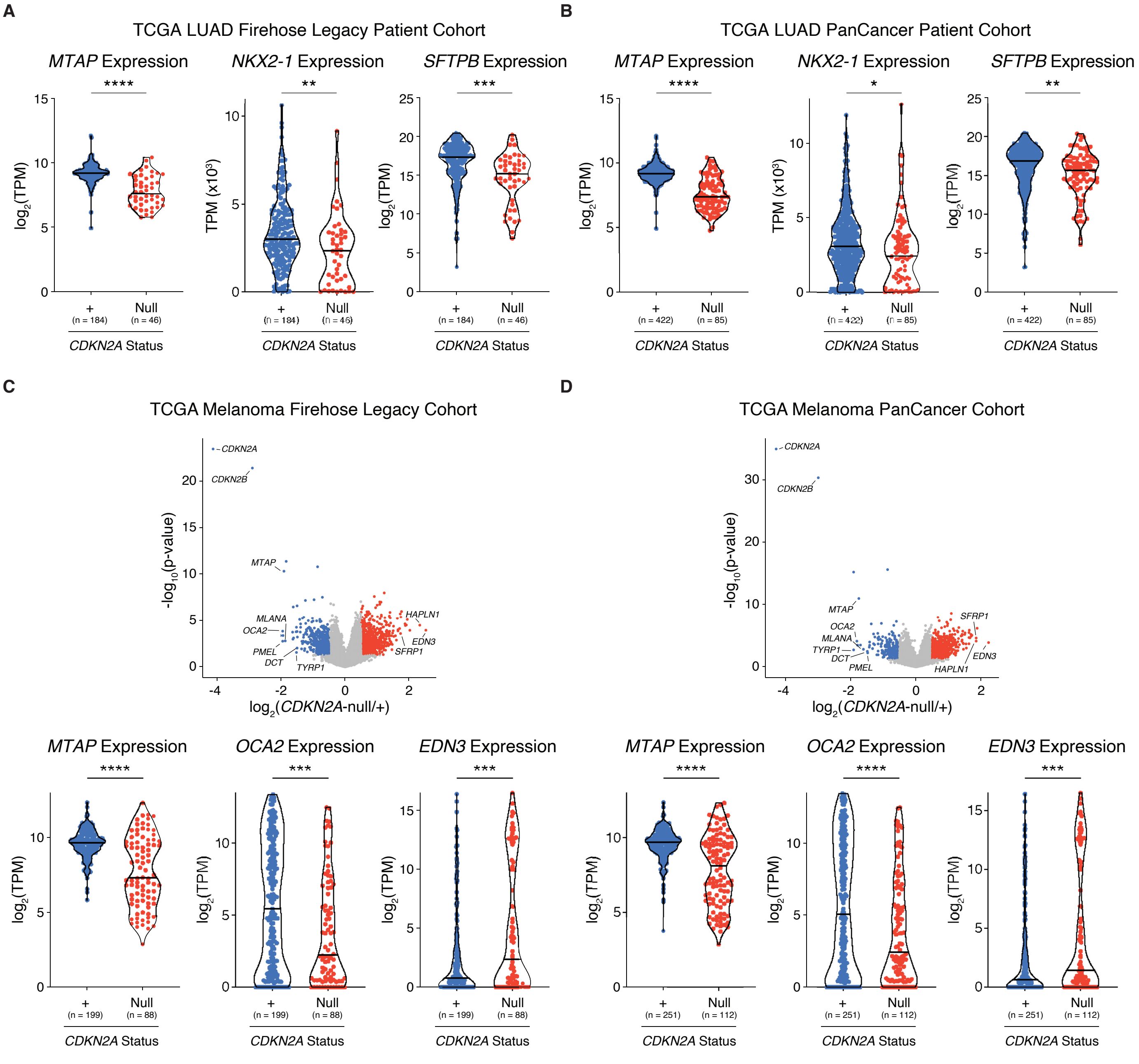

**Supplemental Figure 10. *CDKN2A/MTAP* loss correlates with dedifferentiation in human LUAD and melanoma.**

**A–B.** Expression of *MTAP* and the early-stage lung markers *NKX2-1* and *SFTPB* in *CDKN2A*+ and *CDKN2A*-null tumors from the TCGA LUAD (**A**) Firehose Legacy or (**B**) PanCancer cohorts. Points represent individual patients with the total number of patients per group listed below the axis.

**C–D.** Differentially expressed genes comparing *CDKN2A*-null to the rest of the tumors (*CDKN2A*+) in the TCGA melanoma (**C**) Firehose Legacy and (**D**) PanCancer cohorts. (*Top*) Points represent individual genes that have increased expression (red, $p<0.05$, $log_2FC>0.5$) or decreased expression (blue, $p<0.05$, $log_2FC<-0.5$). (*Bottom*) Expression of *MTAP*, the early-stage melanoma marker *OCA2*, and the late-stage melanoma marker *EDN3* in the indicated groups. Points represent individual patients with the total number of patients per group listed below the axis.

For corresponding panels, Welch's t-test was used. * indicates $p<0.05$, ** indicates $p<0.01$, *** indicates $p<0.001$, and **** indicates $p<0.0001$.
